# THE CONSEQUENCES OF PARTIAL CLONALITY ON THE SPATIOTEMPORAL GENETIC STRUCTURE OF THE RED ALGA *CHONDRIA TUMULOSA* (CERAMIALES, RHODOPHYTA)

**DOI:** 10.1101/2025.09.15.676347

**Authors:** Taylor M. Williams, Heather L. Spalding, Solenn Stoeckel, Brian B. Hauk, Jon Plissner, Randall K. Kosaki, Stacy A. Krueger-Hadfield

**Author notes:** Author for correspondence: Stacy A. Krueger-Hadfield.

## Abstract

The relative rates of sexual versus clonal reproduction partition genetic diversity but the quantification of these rates remains limited in natural populations. This knowledge gap is particularly acute among tropical macroalgae. *Chondria tumulosa* is a red macroalga that has been documented at high abundance in the three northernmost atolls in the Papahānaumokuākea Marine National Monument. We sampled thalli at Manawai in 2021 and 2023, at Kuaihelani in 2022 and 2023, and at Hōlanikū in 2023 to characterize the reproductive mode and patterns of gene flow. Tetrasporophytes overwhelmingly dominated at all sites, atolls, and years, although eight cystocarpic female gametophytes (<1%) were documented for the first time. We observed evidence of very high clonal rates, accompanied by an increase in homozygosity. Surprisingly, our temporal analyses suggested site-specific trends in the predominance of selfing, outcrossing, or clonality. A very small haploid proportion combined with high genetic drift likely rendered temporal analyses of genotypic frequencies challenging to implement and interpret whereby point estimates were a better reflection of reproductive mode variation. Genetic differentiation increased with geographic distance, with strong temporal and spatial structure. The patterns of genetic diversity and structure combined with the dominance of tetrasporophytes suggest that *C. tumulosa* thalli were introduced to this region, possibly via marine debris. Future studies should focus on identifying the native range to document the ecological and evolutionary shifts that have facilitated the rapid increase and spread of *C. tumulosa* and its potential similarities to other invasive haploid-diploid algae.

## INTRODUCTION

Meiotic sex (*sensu* Beukeboom & Perrin, 2014) is responsible for generating tremendous life cycle variation across eukaryotes (Bell, 1994), but our understanding of reproductive mode variation tends to focus on populations of animals (Lane et al., 2011; Olsen et al., 2020) and angiosperms (Eckert et al., 2010; Meirmans et al., 2012, Rushworth et al., 2020). Existing data are also largely based on the relative rates of self-fertilization (hereafter referred to as selfing) versus outcrossing (Barrett et al., 2014). However, a recent review has highlighted a bias in the literature that suggests selfing and mixed mating may be overestimated in angiosperms (Meyer et al., 2024). Although most eukaryotes are partially clonal (Schön et al., 2009), engaging in both sexual and clonal (also called asexual) reproduction, we have largely ignored the contributions of clonal processes in natural populations (Arnaud-Haond et al., 2007; Halkett et al., 2005; Orive & Krueger-Hadfield, 2021). The relative rates of sexual versus clonal reproduction are nevertheless important to discern because of it what partitions genetic diversity within and among populations (Hamrick and Godt, 1996).

Uniparental reproduction presents advantages for populations expanding into new habitats (Pannell et al., 2015). Selfing enables individuals to reproduce even when mates are scarce, thus facilitating population establishment and reproductive assurance during colonization (Razanajatovo et al., 2016). It also facilitates the emergence of gene combinations that may enhance individual fitness in the new environments through recombination between pioneering individuals (Busch & Delph, 2012). Clonal reproduction allows expanding populations to rapidly establish from single individuals, temporally bypassing the need for mates before compatible and unrelated new individuals arrive (Pannell et al., 2015). Clonality also ensures the transmission without recombination of successful genotypes well-suited to the invaded environment (He et al., 2024) and reduces the demographic and genetic costs associated with sexual reproduction, facilitating persistence in new habitats (Pierre et al., 2022).

Empirical data on reproductive system variability is particularly lacking across the phylogenetically diverse lineages of algae (Krueger-Hadfield, 2024). There is anecdotal evidence that macroalgae have substantial reproductive mode variation (see Maggs, 1988; Otto & Marks, 1996; Valero et al., 2001); this has been recently supported by new population genetic data (see reviews in Olsen et al., 2020; Krueger-Hadfield et al., 2024; Krueger-Hadfield, 2024). Even if we can observe reproductive structures, we cannot determine the prevailing reproductive mode in a macroalgal population without population genetics. For example, microsatellite genotyping in Baltic populations of *Fucus vesiculosus* revealed the role of clonal processes in persistence and dispersal (Tatarenkov et al., 2005). Similarly, microsatellite loci were used to distinguish between gametophytes and sporophytes in *Cladophoropsis membranacea*, whereby mats of coalesced thalli were composed of many unique genotypes (van der Strate et al., 2002). Krueger-Hadfield et al. (2021a) re-analyzed data for the 11 haploid-diploid macroalgaefor which gametophytic and sporophytic genotypic data were available, including *C. membrancea*, and demonstrated ample variation in reproductive mode when standardizing analytical methods (Stoeckel et al., 2021). Importantly, Krueger-Hadfield et al. (2021a) and Stoeckel et al. (2021) provided guidelines for which summary statistics to calculate (e.g., *pareto ß,* the distribution of clonal membership) and how to report point estimates of reproductive mode variation (e.g., single locus *F_IS_* and the variance) for haploid-diploid taxa.

The small number of haploid-diploid algae studied using population genetic tools, as reviewed in Olsen et al. (2020) or Krueger-Hadfield et al. (2021a), were focused on temperate macroalgae, with studies in the tropics largely absent. This is true for macroalgae in the Hawaiian Archipelago, despite its rich phycological history (Magruder & Hunt, 1979; Abbott, 1999; Abbott & Huisman, 2004; Tsuda 2014; McDermid et al., 2019; Sherwood & Guiry, 2023). Recently, Thornton et al. (2024) used temporal sampling to demonstrate that clonal processes were responsible for the spread of the green macroalga *Avrainvillea lacerata* at two sites on O ahu, confirming ecological observations by Smith et al. (2002). Similarly, Williams et al. (2024) found genetic signatures of clonality, including repeated genotypes and tetrasporophytic bias, in the red macroalga *Chondria tumulosa*. This alga has been documented at the three most northern atolls in the Northwestern Hawaiian Islands in the Papahānaumokuākea Marine National Monument (PMNM; Sherwood et al., 2020), one of the most remote island archipelagos in the world, covering 1,500,000 km^2^ of the Pacific Ocean (Figure 1; Rohmann et al., 2005). The PMNM has been known as a pristine and intact ecosystem (Maragos & Gulko, 2002; Wilkinson, 2004) and was designated as a United Nations Educational, Scientific, and Cultural Organization (UNESCO) World Heritage Site (Kikiloi et al., 2017) and a National Marine Sanctuary. Yet, coral reefs in the PMNM have not escaped extensive coral bleaching, nor the consequences of marine debris which smother reefs and are a vector for species introductions (PMNM, 2020). The arrival of *C. tumulosa*, coupled with its large biomass (Sherwood et al., 2020) and the genetic signatures reported by Williams et al. (2024), strongly suggest that this alga was introduced to the PMNM and has been categorized as non-native by other authors (Carlton & Schwindt, 2024).

**Figure 1.**
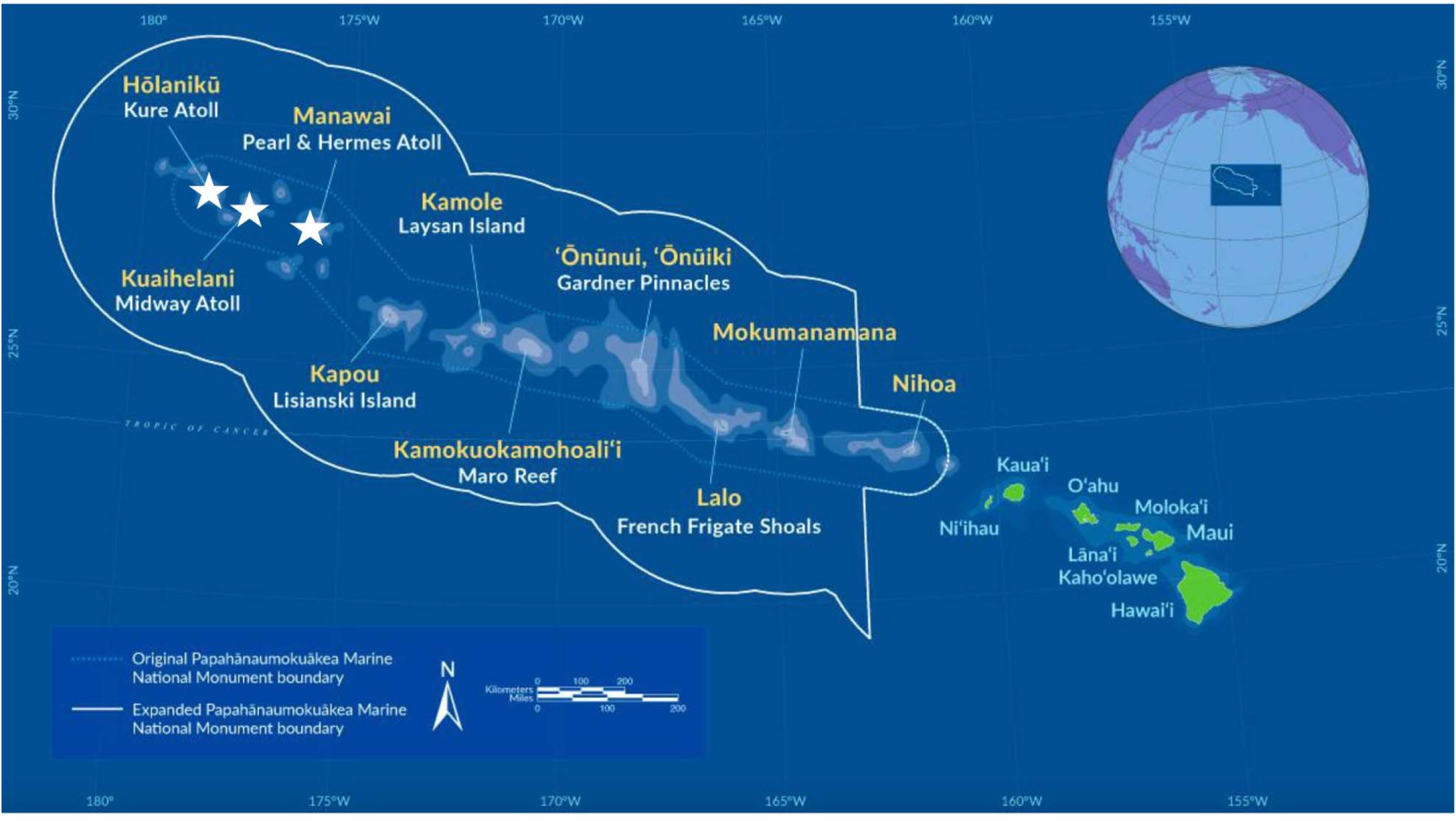
Map of Papahānaumokuākea Marine National Monument (PMNM). The white border shows the boundary line for PMNM. The white stars show the three atolls (Manawai, Kuaihelani, and Hōlanikū) where *Chondria tumulosa* has been identified and collected. Base image courtesy of NOAA PMNM.

*Chondria tumulosa* was first discovered at Manawai (Pearl and Hermes Atoll) in 2015 in very low abundance (Sherwood et al., 2020). By 2019, this alga was smothering coral reefs at Manawai, outcompeting and overgrowing native coral and macroalgal species (Sherwood et al., 2020; Lopes et al., 2023; Williams et al., 2024). Subsequently, it was found at Kuaihelani (Midway Atoll; Lopes et al., 2023) and Hōlanikū (Kure Atoll; this study; Figure 1). There are no data thus far investigating the patterns of genetic diversity and connectivity of *Chondria tumulosa* in the PMNM (but see a modeling approach using particle tracking, Fumo et al., 2024).

Despite genetic signatures of clonal reproduction via thallus fragmentation in 2019, Williams et al. (2024) found some reproductive tetrasporophytes in low abundance, although the fate of the tetraspores is unknown. For most marine haploid-diploid red algae in the Florideophyceae, sexual reproduction involves the production of gametes and spores, subsequent recruitment, and eventual reproductive maturity of two independent phases – the gametophytes and tetrasporophytes. In 2019, only tetrasporophytes were observed, based on the presence of tetrasporangial sori, heterozygous genotypes from microsatellite genotyping, or both (Williams et al., 2024). Using nine polymorphic microsatellite loci, we expanded the study conducted in 2019 and describe the reproductive mode using both single time point and temporal sampling to quantitatively infer the respective rates of clonality, outcrossing, and selfing. Based on previous observations (Williams et al., 2024) and the fact that *C. tumulosa* may benefit from the ecological and genetic advantages of clonal reproduction during expansion (e.g., Krueger-Hadfield et al., 2016; 2020), we predicted that clonality through thallus fragmentation would be the prevailing reproductive mode, resulting in stable tetrasporophytic bias across all atolls and time points. We also investigated patterns of genetic structure over space and time to understand the potential for *C. tumulosa* spreading in the PMNM. We expected to find evidence of limited gene flow between atolls and directionality from Manawai to the other two northernmost atolls based on the possible expansion front. Together, these data are critical for understanding connectivity across these three atolls and the dispersal mechanisms that may spread *C. tumulosa* throughout the PMNM and to the Main Hawaiian Islands.

## METHODS

### Sample collection

*Chondria tumulosa* thalli were collected as small, palm-sized clumps at sites located at three atolls in PMNM, which from 2019 to 2023 encompassed all known locations of this macroalga (Figure 2, Figure 3, Table 1): (1) Manawai in July 2019 (N = 5 sites, N = 124 clumps; see also Williams et al., 2024), July 2021 (N = 8 sites, N = 499 samples) and July 2023 (N = 10 sites, N = 549 samples); (2) Kuaihelani in July 2022 (N = 6 sites, N = 274 samples), October 2022 (only site MID_33, N = 30 samples), and July 2023 (only MID_33, N = 50 samples); and (3) Hōlanikū in July 2023 (N =4 sites, N = 103 samples). Clumps were haphazardly collected using SCUBA approximately every meter to prevent collecting from the same area twice.

**Figure 2.**
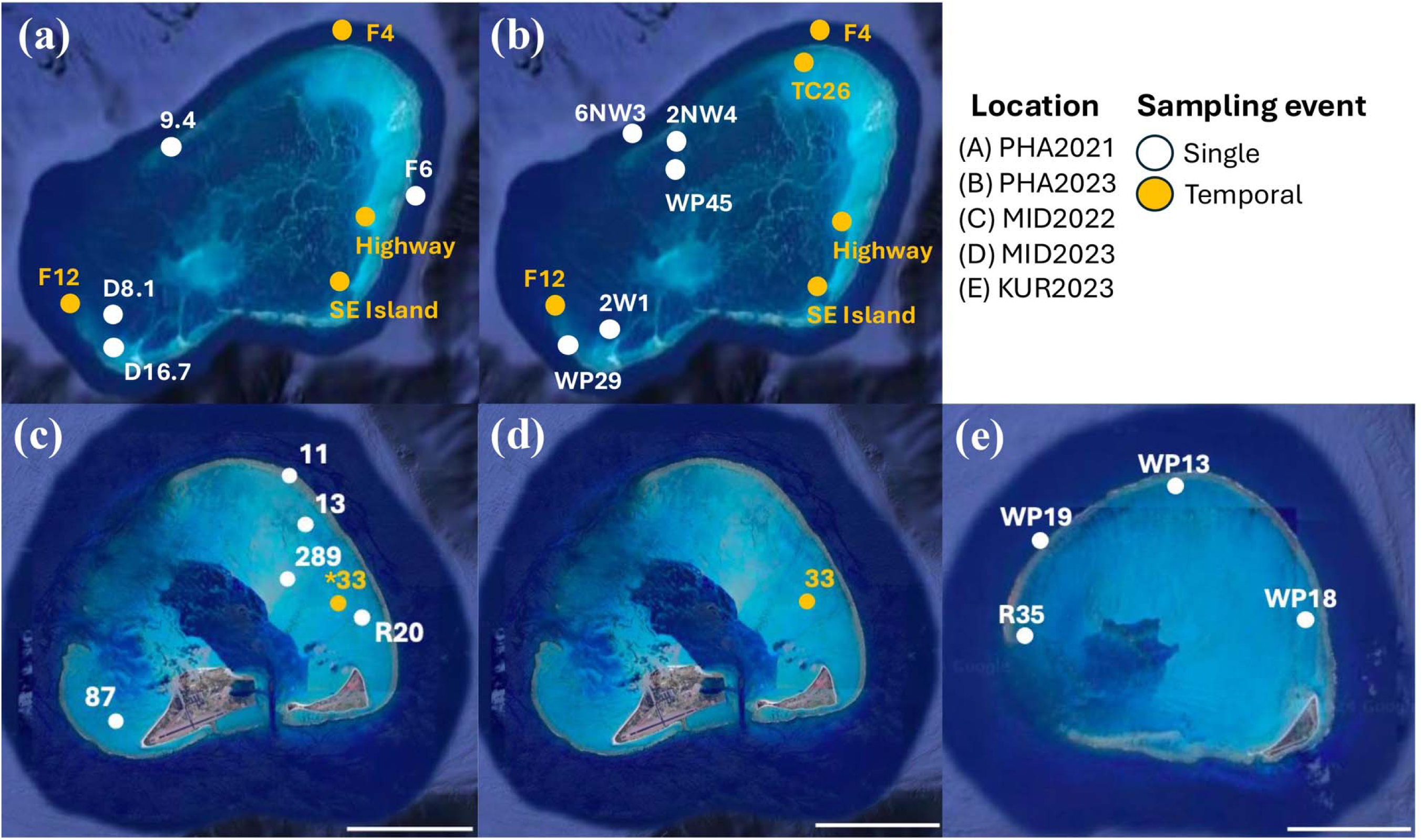
Sampling locations at Manawai (PHA2021, PHA2023), Kuaihelani (MID2022, MID2023), and Hōlanikū (KUR2023). Scale bars indicate 1.5 kilometers. *MID_33 was sampled in July and October 2022. Base image of atolls from Google Maps 2024.

**Figure 3.**
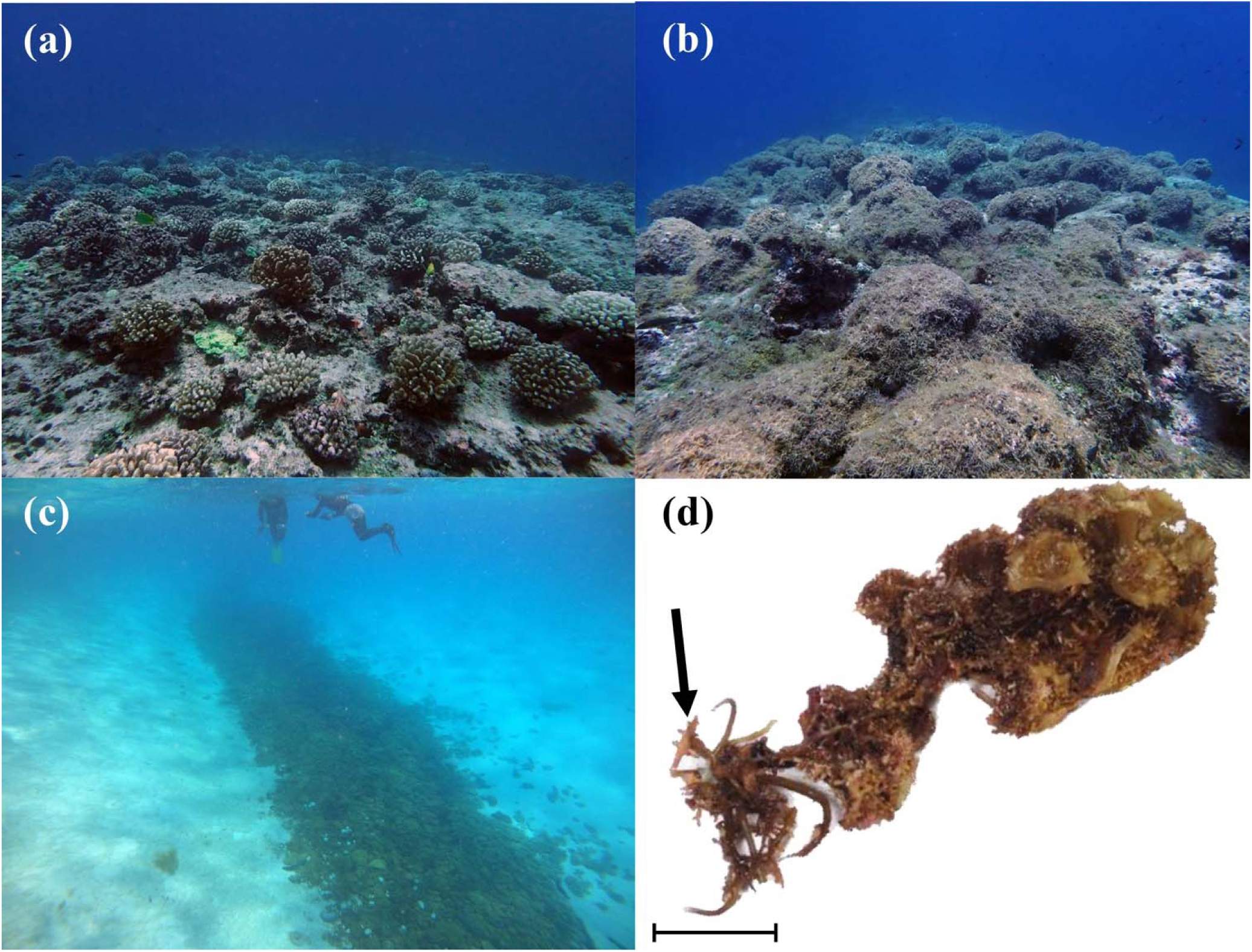
Representative photos of the habitats in which *Chondria tumulosa* is now found in the Papahānaumokuākea Marine National Monument (PMNM). (A) A site at Manawai with low percent cover of *C. tumulosa* on a *Pocillopora meandrina* coral reef (photo credit: H.L. Spalding). (B) A site at Manawai with high percent cover of *C. tumulosa* on a *P. meandrina* coral reef (photo credit: T.M. Williams). (C) Free-living *C. tumulosa* thalli accumulating in sandy lagoon at Manawai (photo credit: B.B. Hauk). (D) At Kuaihelani, *C. tumulosa* thalli were found growing epiphytically on *Turbinaria ornata*. The black arrow points to representative *C. tumulosa* thallus (photo credit: T.M. Williams). Scale bar = 2 cm.

The clumps of *Chondria tumulosa* thalli collected from mats were previously found to contain more than one genotype (Williams et al., 2024). Therefore, we selected representative thalli (2 to 4 thalli depending on variability in thallus morphology) from each clump and scanned them using a 4X objective with 10X oculars on a compound microscope (Leica DM500, Deerfield, Illinois, USA) to determine the reproductive state of the fragment (N = 4,027). If tetrasporangial sori were observed, then the thallus was considered a tetrasporophyte. If spermatangial sori were observed, then the thallus was considered a male gametophyte. If carposporophytes were observed, the thallus was considered a female gametophyte. Finally, if there were no observable reproductive structures, the thallus was considered vegetative at the time of sampling and the phase (and sex if applicable) was not determined. We note that it would have been difficult to observe carpogonia prior to fertilization or trichogynes on a female gametophyte, and therefore some of the vegetative thalli may have been reproductive but unfertilized females. A representative thallus was then selected from those above to be preserved in silica (ACTIVA Art Silica Gel, Marshall, Texas, Cat No. NOTM691267) for subsequent DNA extraction.

### Fragment analysis and genotyping

Total genomic DNA was isolated using a Nucleospin^®^ 96 Plant Kit (Macherey-Nagel, Düren, Germany) from 1,174 thalli (Table 1A). The manufacturer’s instructions were followed for all steps except the lysis step, in which the lysate was held at room temperature for one hour, and the final elution, which was performed in 100 µl of molecular grade water (see Krueger-Hadfield et al., 2013, as well as Williams et al., 2024).

**Table 1A.** Metadata and sample size for sites from which we collected *Chondria tumulosa* thalli. Site name, atoll abbreviation (PHA, Manawai; MID, Kuaihelani; KUR, Hōlanikū) and site name; Site no., each site was assigned a number for ease of visualization on plots; Year, sampling year; Latitude; Longitude; Depth, sampling depth in meters; Mat forming, yes or no as to whether the site had *C. tumulosa* mats; Percent cover, amount of *C. tumulosa* cover at the site; No. collected, the number of clumps of *Chondria tumulosa* thalli collected at the site; No. repro. State, the number of *C. tumulosa* thalli that were examined for reproductive structures; No. genotyped, the number of *C. tumulosa* thalli genotyped at nine microsatellite loci; No. with genotypes, the number of successfully genotyped *C. tumulosa* thalli used in downstream analyses. Sites sampled at more than one time point are assigned the same site number and are shown in bold with gray shading.

| Site name | Site no. | Year | Latitude | Longitude | Depth | Mat forming? | Percent cover | No. clumps | No. repro. state | No. genotyped | No. with genotypes |
| --- | --- | --- | --- | --- | --- | --- | --- | --- | --- | --- | --- |
| PHA_16.7 | 6 | 2021 | 27.7902 | -176.0001 | 17 | No | 40 | 50 | 150 | 50 | 50 |
| PHA_D8.1 | 7 | 2021 | 27.7904 | -175.9962 | 9 | Yes | 75 | 50 | 150 | 50 | 50 |
| PHA_F6 | 8 | 2021 | 27.8515 | -175.7402 | 13 | No | 55 | 49 | 147 | 49 | 49 |
| **PHA_F4** | **9** | **2021** | **27.9637** | **-175.7742** | **13** | **No** | **50** | **50** | **150** | **50** | **50** |
| PHA_9.4 | 10 | 2021 | 27.8981 | -175.9234 | 9 | Yes | 70 | 100 | 300 | 100 | 99 |
| **PHA_F12** | **11** | **2021** | **27.8185** | **-175.9883** | **14** | **No** | **45** | **50** | **150** | **50** | **50** |
| **PHA_Highway** | **12** | **2021** | **27.8185** | **-175.7628** | **5** | **No** | **100** | **100** | **300** | **100** | **100** |
| **PHA_SEIsland** | **13** | **2021** | **27.7953** | **-175.8207** | **7** | **No** | **70** | **50** | **150** | **50** | **50** |
| PHA_WP029 | 15 | 2023 | 27.7897 | -175.9953 | 13 | No | 10 | 50 | 99 | 25 | 23 |
| PHA_WP045 | 16 | 2023 | 27.9069 | -175.8884 | 3 | No | 10 | 50 | 100 | 25 | 23 |
| **PHA_F4** | **9** | **2023** | **27.9638** | **-175.7741** | **14** | **No** | **50** | **50** | **100** | **26** | **26** |
| PHA_TC26 | 17 | 2023 | 27.9578 | -175.8021 | 2 | Yes | 100 | 50 | 100 | 25 | 25 |
| **PHA_SEIsland** | **13** | **2023** | **27.8304** | **-175.8021** | **1** | **No** | **70** | **50** | **100** | **25** | **23** |
| **PHA_Highway** | **12** | **2023** | **27.8850** | **-175.7572** | **4** | **No** | **100** | **50** | **100** | **25** | **24** |
| PHA_2W1 | 18 | 2023 | 27.8122 | -175.9960 | 15 | Yes | 80 | 50 | 100 | 25 | 24 |
| **PHA_F12** | **11** | **2023** | **27.8185** | **-175.9879** | **11** | **No** | **45** | **50** | **100** | **25** | **24** |
| PHA_2NW4 | 19 | 2023 | 27.9257 | -175.8832 | 14 | No | 20 | 50 | 100 | 25 | 24 |
| PHA_6NW3 | 20 | 2023 | 27.9101 | -175.9033 | 9 | Yes | 80 | 50 | 100 | 25 | 24 |
| MID_11 | 21 | 2022 | 28.2702 | -177.3411 | 2 | No | 20 | 49 | 145 | 49 | 50 |
| MID_13 | 22 | 2022 | 28.2577 | -177.3345 | 2 | No | 10 | 50 | 150 | 50 | 49 |
| **MID_33** | **23** | **2022July** | **28.2431** | **-177.3372** | **3** | **Yes** | **80** | **50** | **150** | **50** | **50** |
| **MID_33** | **23** | **2022Oct** | **28.2431** | **-177.3372** | **3** | **Yes** | **80** | **30** | **0** | **30** | **30** |
| MID_87 | 24 | 2022 | 28.2009 | -177.4096 | 1 | Yes | 80 | 50 | 150 | 50 | 50 |
| MID_WP289 | 25 | 2022 | 28.2478 | -177.3530 | 1 | Yes | 100 | 50 | 150 | 50 | 50 |
| MID_R20 | 26 | 2022 | 28.231 | -177.3291 | 3 | No | 5 | 25 | 75 | 25 | 24 |
| **MID_33** | **23** | **2023** | **28.2432** | **-177.3373** | **2** | **Yes** | **80** | **50** | **100** | **50** | **50** |
| KUR_R35 | 27 | 2023 | 28.4197 | -178.3719 | 5 | No | 1 | 20 | 36 | 20 | 19 |
| KUR_WP013 | 28 | 2023 | 28.4535 | -178.3286 | 2 | No | 1 | 17 | 18 | 17 | 9 |
| KUR_WP018 | 29 | 2023 | 28.4215 | -178.2889 | 2 | No | 1 | 24 | 30 | 24 | 14 |
| **KUR_WP019** | **30** | **2023** | **28.4407** | **-178.3608** | **15** | **No** | **1** | **24** | **38** | **24** | **19** |
| Total | – | – | – | – | – | – | – | 1,438 | 3,538 | 1,189 | 1,152 |

**Table 1B.** Summary of counts in Dataset One, including all reproductive tetrasporophytes, all heterozygous thalli, and all repeated multilocus genotypes that were identical to a reproductive tetrasporophyte (including fixed homozygous thalli). N, total sample size of thalli genotyped; Repro. tetra, number of reproductive tetrasporophytes based on microscopy that were later genotyped; Repro. gameto, number of reproductive gametophytes based on microscopy that were later genotyped; Vegetative thalli, number of vegetative thalli based on microscopy that were later genotyped; Heterozyg, thalli, number of heterozygous thalli based on 9 microsatellite loci; Homozyg. thalli, number of homozygous thalli based on 9 microsatellite loci; Total tetra., the total number of tetrasporophytes based on microscopy and microsatellite genotyping; Total gameto., the total number of gametophytes based on microscopy and microsatellite genotyping; *P_HD_*, ploidy diversity (* indicates significant deviation from the predicted phase ratio of √2:1, see Table S3). Sites sampled more than once are shown in bold and gray shading.

| Site name | Site No. | Year | N | Repro. tetra. | Repro gameto | Vegetative thalli | Heterozyg. thalli | Homozyg. thalli | Total tetra. | Total gameto. | $P_{HD}$ |
| --- | --- | --- | --- | --- | --- | --- | --- | --- | --- | --- | --- |
| PHA_16.7 | 6 | 2021 | 50 | 5 | 0 | 45 | 44 | 4 | 44 | 6 | 0.20* |
| PHA_D8.1 | 7 | 2021 | 50 | 21 | 0 | 29 | 42 | 2 | 43 | 7 | 0.24* |
| PHA_F6 | 8 | 2021 | 49 | 5 | 0 | 44 | 44 | 4 | 44 | 5 | 0.17* |
| **PHA_F4** | **9** | **2021** | **50** | **12** | **1** | **37** | **40** | **8** | **44** | **6** | **0.20\*** |
| PHA_9.4 | 10 | 2021 | 99 | 1 | 0 | 98 | 56 | 40 | 56 | 43 | 0.74* |
| **PHA_F12** | **11** | **2021** | **50** | **14** | **0** | **36** | **48** | **1** | **48** | **2** | **0.07\*** |
| **PHA_Highway** | **12** | **2021** | **100** | **19** | **0** | **81** | **88** | **8** | **89** | **11** | **0.19\*** |
| **PHA_SEIsland** | **13** | **2021** | **50** | **11** | **1** | **38** | **50** | **0** | **50** | **0** | **0.00\*** |
| PHA_WP029 | 15 | 2023 | 23 | 5 | 0 | 18 | 20 | 0 | 20 | 3 | 0.22* |
| PHA_WP045 | 16 | 2023 | 23 | 0 | 0 | 23 | 21 | 2 | 21 | 2 | 0.15* |
| **PHA_F4** | **9** | **2023** | **26** | **1** | **0** | **25** | **21** | **5** | **22** | **4** | **0.26\*** |
| PHA_TC26 | 17 | 2023 | 25 | 0 | 0 | 25 | 2 | 21 | 3 | 23 | 0.20* |
| **PHA_SEIsland** | **13** | **2023** | **23** | **3** | **0** | **20** | **22** | **0** | **22** | **1** | **0.07\*** |
| **PHA_Highway** | **12** | **2023** | **24** | **4** | **0** | **20** | **24** | **0** | **24** | **0** | **0.00\*** |
| PHA_2W1 | 18 | 2023 | 24 | 6 | 0 | 18 | 22 | 1 | 22 | 2 | 0.14* |
| **PHA_F12** | **11** | **2023** | **24** | **6** | **0** | **18** | **21** | **2** | **22** | **2** | **0.14\*** |
| PHA_2NW4 | 19 | 2023 | 24 | 4 | 0 | 20 | 19 | 3 | 20 | 4 | 0.28* |
| PHA_6NW3 | 20 | 2023 | 24 | 2 | 0 | 22 | 14 | 10 | 14 | 10 | 0.71* |
| MID_11 | 21 | 2022 | 50 | 0 | 0 | 50 | 30 | 19 | 30 | 20 | 0.68* |
| MID_13 | 22 | 2022 | 49 | 3 | 0 | 46 | 37 | 6 | 37 | 12 | 0.42* |
| **MID_33** | **23** | **2022** | **50** | **5** | **0** | **45** | **32** | **15** | **33** | **17** | **0.58\*** |
| MID_87 | 24 | 2022 | 50 | 3 | 0 | 47 | 40 | 10 | 42 | 8 | 0.27* |
| MID_WP289 | 25 | 2022 | 50 | 20 | 0 | 30 | 9 | 39 | 14 | 36 | 0.68* |
| MID_R20 | 26 | 2022 | 24 | 2 | 1 | 22 | 10 | 13 | 12 | 12 | 0.85 |
| **MID_33** | **23** | **2023** | **50** | **4** | **0** | **45** | **34** | **15** | **35** | **15** | **0.51\*** |
| KUR_R35 | 27 | 2023 | 19 | 2 | 0 | 17 | 10 | 8 | 10 | 9 | 0.80 |
| KUR_WP013 | 28 | 2023 | 9 | 2 | 0 | 7 | 1 | 8 | 2 | 7 | 0.54 |
| KUR_WP018 | 29 | 2023 | 14 | 3 | 1 | 11 | 1 | 13 | 4 | 10 | 0.70 |
| KUR_WP019 | 30 | 2023 | 19 | 11 | 0 | 8 | 6 | 12 | 10 | 9 | 0.80 |

We used five microsatellite loci from Williams et al. (2024) and added an additional four loci (Table S1). Polymerase Chain Reactions (PCRs) were performed to amplify nine loci in simplex using a SimpliAmp™ thermocycler (ThermoFisher Scientific, Waltham, MA) or a PROFLEX^®^ thermocycler (Proflex, Haines City, Florida). The following PCR program was used: 2 min at 95°C, followed by 30 cycles of 95°C for 30 sec, 56°C for 30 sec, and 72°C for 30 sec, with a final extension of 5 min at 72°C. The PCR final volume of 10 µL contained 2 µl DNA template, 1X Promega buffer, 250 µM of each dNTP, 2 mM MgCl_2_, 1 mg . ml^−1^ bovine serum albumin (BSA), 1 U Promega GoFlexi *Taq* polymerase, 250 nM fluorescently labeled forward primer (using 6-FAM, VIC, NED, or PET; see Table S1), 150 nM unlabeled forward primer, and 400 nM unlabeled reverse primer. One µL of PCR product was diluted in 9.7 µL of Hi-Di Formamide (ThermoFisher Scientific) and 0.35 µL size ladder (GeneScan^TM^ 500 LIZ^TM^ dye Size Standard, ThermoFisher Scientific) for fragment analysis at the University of Alabama at Birmingham Heflin Center for Genomic Sciences (Birmingham, Alabama, USA). Poolplex pools were as follows: *Chondria*_10, *Chondria*_34, *Chondria*_74, and *Chondria*_44 pooled together and *Chondria*_39, *Chondria*_29, *Chondria*_24, and *Chondria*_7 pooled together.

Alleles were scored manually using GENEIOUS PRIME v. 2020.0.5 (Biomatters, Ltd., Auckland, New Zealand). For the new loci *Chondria*_10, *Chondria*_44, *Chondria*_70, and *Chondria*_74, we used TANDEM (Matschiner and Salzburger, 2009) to bin alleles. For the other five loci, alleles were binned manually based on allele calls defined in Williams et al. (2024). We binned alleles and created a scoring guide for future use of these loci (Table S2).

### Phase determination and ratio

Most thalli appeared vegetative at the time of collection. Thus, the phase (haploid gametophyte vs. diploid tetrasporophyte) was assigned using the multilocus genotype (MLGs) following Krueger-Hadfield et al. (2013) and using the methods implemented in Krueger-Hadfield et al. (2016). Fragment analysis software often assumes diploidy, and two alleles are therefore assumed for each locus. All reproductive gametophytes appeared as fixed homozygotes (see also Kollars et al., 2015; Heiser et al., 2023a; Crowell et al., 2024). If at least one locus was heterozygous, the thallus was considered a diploid tetrasporophyte. If all loci were homozygous but the thallus was a reproductive tetrasporophyte at the time of collection, the thallus was considered a fixed homozygous tetrasporophyte (see Krueger-Hadfield et al., 2013 or Heiser et al., 2023b for more discussion on fixed homozygous tetrasporophytes). If all loci were homozygous and the thallus was vegetative at the time of collection, we currently have no other method with which to determine phase, such as a chemical test (e.g., the resorcinol test, Lazo et al., 1989) or sex-linked genetic markers (e.g., Krueger-Hadfield et al., 2021b). Thus, we considered two different datasets for the summary statistics described in the following subsections: Dataset One included (i) all reproductive tetrasporophytes, (ii) all heterozygous thalli, and (iii) any fixed homozygous thalli that matched at all loci to a reproductive tetrasporophyte (i.e., repeated multilocus genotype, MLG); and Dataset Two included (i) all reproductive tetrasporophytes and (ii) all heterozygous thalli, but we treated any vegetative, fixed homozygous thallus as a haploid gametophyte as has been done previously in other red algae (see Krueger-Hadfield et al., 2016; Heiser et al., 2023b). We focused all our population genetic analyses on the tetrasporophytes at each site because of the tetrasporophytic bias in both datasets (see also Krueger-Hadfield et al., 2016; Heiser et al., 2023b).

We compared the two datasets to ensure patterns were consistent using Wilcoxon signed rank tests or paired t-tests in R ver. 4.3.3 (R Core Team, 2024) depending on whether each summary statistic met the assumptions of normality. We only report the results from Dataset One in the main text, with data from Dataset Two provided in supplementary information. Dataset One, although less conservative, better represents the tetrasporophytic bias evident at all sites sampled thus far and is consistent with the ecology of this species in the PMNM (see also Williams et al., 2024).

Deviations from the predicted ploidy ratio of √2:1 (0.41 tetrasporophytes to 0.59 gametophytes) were detected using the binomial distribution (Destombe et al., 1989; Thornber and Gaines, 2004). We also calculated ploidy diversity (*P_HD_*) following Krueger-Hadfield et al. (2019).

### Genotypic and genetic diversity

We followed the suggestions of Krueger-Hadfield et al. (2021a) and Stoeckel et al. (2021) for analyzing population genetic data in partially clonal, haploid-diploid taxa. First, to determine the resolution of our suite of microsatellite loci, we calculated the unbiased probability of identity (*pidu*) and the probability of identity between sibs (*pids*) (Waits et al., 2010). Null allele frequencies were calculated using ML-Null Freq (Kalinowski & Taper, 2006) in the tetrasporophytes. Null allele frequency in the gametophytes was calculated as the number of loci with non-amplification after discounting technical errors (see Krueger-Hadfield et al., 2013).

We calculated genotypic richness as 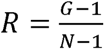, with *G* being the number of distinct genotypes and *N* as the total number of genotyped thalli (Dorken and Eckert, 2001). We also calculated genotypic evenness (*D\**) to describe the distribution of sampling units among clonal lineages (Arnaud-Haond et al., 2007). For genotypic evenness, if a site is dominated by a few dominant clones, *D\** approaches 0. In contrast, if each genet is represented by an equal number of ramets at a site, *D\** will approach 1 even if there are repeated MLGs. To facilitate interpretation, we explored the relationship between genotypic richness and genotypic evenness following Baums et al. (2006) and Krueger-Hadfield et al. (2021a). For each MLG we computed *psex*, or the probability of encountering the MLG’s genotype more than once by chance considering the population allele frequencies (Arnaud-Haond et al., 2007, Bailleul et al., 2016).

Next, we calculated *pareto ß* to estimate the distribution of clonal membership (Arnaud-Haond et al., 2007). Krueger-Hadfield et al. (2021a) used available data at the time to set threshold *pareto ß* values for clonal rates and we used these same ranges: a value < 0.7 suggests high clonal rates, values between 0.7 and 2.0 suggest partial clonality, and values > 2.0 suggest low rates of clonality (Krueger-Hadfield et al., 2021a). Additionally, we calculated a multilocus estimate of linkage disequilibrium (*r̄_D_*) to measure non-random associations between alleles, expected heterozygosity (*H_E_*) and observed heterozygosity (*H_O_*) to assess genetic variation within populations, and the inbreeding coefficient (*F_IS_*) to assess deviations from Hardy-Weinberg proportions in GenAPoPop (Stoeckel et al., 2024a). For *H_E_*, *H_O_*, and *F_IS_*, we also calculated their variance.

To then infer if the reproductive mode could explain the transition of genotype frequencies between each year sampled, we used GenAPoPop (Stoeckel et al., 2024a). We used the uniform joint priors covering the mutation rate u ∈ [0.001], the clonal rate c ∈ [0.0, 0.1, 0.2, 0.3, 0.4, 0.5, 0.6, 0.7, 0.8, 0.9, 1.0], and the selfing rate s ∈ [0.0, 0.1, 0.2, 0.3, 0.4, 0.5, 0.6, 0.7, 0.8, 0.9, 1.0] to cover the full spectrum of all possible quantitative reproductive modes. This was done at sites for which samples had been collected at more than a single time point at Manawai (PHA) and Kuaihelani (MID; Table 1A). At Kuaihelani, the only site that was sampled at more than one timepoint was MID_33, in which we sampled thalli in July 2022, October 2022, and July 2023 and we performed comparisons between July 2022 vs. October 2022, July 2022 vs. July 2023, and October 2022 vs. July 2023. There were four sites at Manawai that were sampled in July 2021 and July 2023: PHA_F12, PHA_F4, PHA_SEIsland, and PHA_Highway. PHA_F12 and PHA_F4 are characterized by thalli that likely re-attach to the substrate (see Heiser et al., 2023b for a discussion on secondary reattachment and Krueger-Hadfield et al., 2023) and percent cover at these sites was around 45% and 50%, respectively (Figure 3). PHA_SE Island and PHA_Highway were both entirely composed of drift thalli in a sandy bottom habitat (Figure 3).

### Ranking sites by clonality and selfing

We used the categorical sums of ranks of population genetic indices known to vary with rates of clonality and the sums of the normalized values of these same indices Σ*clon* to rank the studied sites from the least to the most clonal following Stoeckel et al. (2024b) proposal and recommendations. These normalized ranking indices compute for each population *i* as 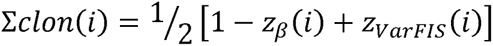 where z indices are the normalized values of each population value over the whole dataset computed as 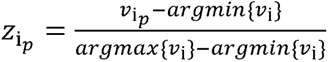 in which (is the considered genetic index, p is the considered population, *argmax*{*v_i_*} is the maximum value of the (genetic index over all studied populations and *argmax*{*v_i_*} is the minimum value of the *i* genetic index over all studied populations. We then tested if sampled sites differ in these relative rankings of population from the less to the more clonal using a Kruskall-Wallis test. We used a Conover post-hoc test with Holm correction to obtain the adjusted p-values between pairwise sampled sites using Scipy and Scikit-posthoc (Virtanen et al., 2020).

### Genetic differentiation

The genetic distance between thalli was then visualized using a minimum spanning tree of the pairwise genetic distance as the number of shared alleles to better interpret genetic ancestry between samples in hierarchically structured populations (Stoeckel et al., 2024b). For the minimum spanning tree, the thalli collected from Manawai in 2019 were also included (see Williams et al., 2024).

We also estimated pairwise genetic differentiation between the sites by calculating *F*_ST_ (Excoffier et al., 1992) using GenoDive ver. 3.06 (Meirmans, 2020). We measured the most direct pathway, which is the geographic distance in kilometers (km) between two sites taking into consideration the contours of the reef on each atoll. This was measured using the measure distance tool in Google^®^ Maps considering the curvature of the Earth’s surface. We then measured geographic distances, which is the most direct distance in kilometers (km) between two sites. We then performed Mantel tests to detect relationships between genetic and direct and shortest-maritime distances between each site using Passage2 (Rosenberg & Anderson, 2011). To test if founder effects would be responsible for initial differentiation that would then decrease over time with migration, we performed Mann-Whitney tests of the *F*_ST_ between sites in Manawai (PHA) over the two sampled years (2021 and 2023). We expect significant decrease if genetic differentiation was only-initially caused by founder effect, while we expect it to remain stable or even increasing if it is due to spatial structure of subpopulations.

### Data Visualization

Figures were prepared using R with the following packages: ggplot2 (Wickham, 2016), ggrepel (Slowikowski, 2024), ggridges (Wilke, 2024), dplyr (Wickham et al., 2022), tidyverse (Wickham et al., 2019), corrplot (Wei & Simko, 2024), and corrr (Kuhn et al., 2022).

## RESULTS

We sampled 1,438 clumps of *C. tumulosa* material (Table 1A). From these samples, we examined 3,538 thalli for reproductive structures and genotyped 1,189 thalli. Of the 1,189 thalli that were genotyped using fragment analysis, only 1,152 were amplified at all loci (see Table 1A). Thirty-seven thalli had poor amplification across all loci and were removed from the dataset.

All new loci had an average rounding error below the recommended error (Table S1) as assessed by TANDEM (Matschiner & Salzburger, 2009). All loci were polymorphic with 2 to 8 alleles at any given locus (Table S2). There were no novel alleles for the five microsatellite loci previously used in Williams et al. (2024). The thalli collected in 2019 were not re-analyzed in this study because too few thalli were successfully genotyped at the additional loci, likely due to low DNA yields following re-extraction of single thalli (see discussion of sampling and preservation issues in Williams et al., 2024).

For the tetrasporophytes, null alleles frequencies calculated using ML-Null Freq ranged from 0% to 15%, apart from the locus Ct_75 with a frequency of 32% at site PHA_TC26 in 2023 (Table S1). All thalli we considered gametophytes amplified at all loci. Therefore, there were no null alleles detected in the gametophytes.

The overall unbiased probability of identity between sibs (*pidu*) was 0.019 and the *pids* was 0.178, suggesting low power to resolve between genotypes probably due to restricted ancestral ties between pioneering individuals. Thus, we interpreted the following patterns with caution (see site-level *pidu* and *pids* values in Table 2).

**Table 2.** Descriptive statistics calculated on Dataset One, including all reproductive tetrasporophytes, all heterozygous thalli, and all repeated multilocus genotypes that were identical to a reproductive tetrasporophyte (including fixed homozygous thalli). N, total sample size of thalli genotyped; *pidu*, unbiased probability of identity of sibs; *pids*, probability of identity of sibs; *G*, number of genotypes; *R*, genotypic richness: *D\**, genotypic evenness; Rep MLG, the number of repeated MLGs (and the range of re-encounters x times); *Pareto ß*, the distribution of clonal membership; *r̄_D_*, multilocus estimate of linkage disequilibrium; *H_E_* (var), multilocus estimate of expected heterozygosity and variance; and *H_O_* (var), multilocus estimate of observed heterozygosity and variance; *F_IS_* (var), multilocus estimate of the inbreeding coefficient and variance. Sites sampled more than once are shown in bold and gray shading.

| Site name | Site No. | Year | N | *pidu* | *pids* | *G* | *R* | *D*<sup>*</sup> | Rep MLG | *Pareto* $\beta$ | $\bar{r}_D$ | *H<sub>E</sub>* (var) | *H<sub>O</sub>* (var) | *F<sub>IS</sub>* (var) |
| --- | --- | --- | --- | --- | --- | --- | --- | --- | --- | --- | --- | --- | --- | --- |
| PHA_16.7 | 6 | 2021 | 50 | 0.011 | 0.145 | 26 | 0.53 | 0.95 | 24 (1–8x) | 0.75 | 0.071 | 0.21 (0.31) | 0.19 (9.53) | 0.06 (3.10) |
| PHA_D8.1 | 7 | 2021 | 50 | 0.015 | 0.184 | 22 | 0.49 | 0.93 | 22 (1–9x) | 0.71 | 0.022 | 0.18 (0.31) | 0.18 (0.37) | 0.01 (3.49) |
| PHA_F6 | 8 | 2021 | 49 | 0.012 | 0.151 | 28 | 0.57 | 0.95 | 28 (1–9x) | 0.80 | 0.019 | 0.20 (0.38) | 0.19 (0.55) | 0.04 (3.05) |
| **PHA_F4** | **9** | **2021** | **50** | **0.011** | **0.153** | **25** | **0.51** | **0.95** | **25 (1–6x)** | **0.86** | **0.040** | **0.20 (0.32)** | **0.19 (0.49)** | **0.04 (3.47)** |
| PHA_9.4 | 10 | 2021 | 99 | 0.041 | 0.330 | 22 | 0.22 | 0.81 | 21 (1–38x) | 0.25 | 0.170 | 0.12 (0.14) | 0.22 (0.11) | 0.25 (1.23) |
| **PHA_F12** | **11** | **2021** | **50** | **0.033** | **0.224** | **14** | **0.27** | **0.86** | **14 (1–12x)** | **0.27** | **0.002** | **0.16 (0.34)** | **0.18 (0.71)** | **-0.12 (7.96)** |
| **PHA_Highway** | **12** | **2021** | **100** | **0.022** | **0.175** | **36** | **0.37** | **0.96** | **32 (1–11x)** | **0.81** | **0.009** | **0.19 (0.38)** | **0.18 (0.43)** | **0.04 (1.84)** |
| **PHA_SEIsland** | **13** | **2021** | **50** | **0.076** | **0.348** | **11** | **0.20** | **0.53** | **11 (1–34x)** | **0.10** | **0.134** | **0.11 (0.19)** | **0.16 (0.81)** | **-0.44 (6.34)** |
| PHA_WP029 | 15 | 2023 | 23 | 0.051 | 0.275 | 10 | 0.47 | 0.91 | 10 (1–5x) | 0.90 | -0.015 | 0.14 (0.35) | 0.16 (0.44) | -0.13 (7.66) |
| PHA_WP045 | 16 | 2023 | 23 | 0.042 | 0.263 | 11 | 0.45 | 0.91 | 11 (1–6x) | 0.76 | -0.015 | 0.14 (0.30) | 0.15 (0.46) | -0.08 (5.97) |
| **PHA_F4** | **9** | **2023** | **26** | **0.022** | **0.205** | **16** | **0.60** | **0.94** | **15 (1–6x)** | **0.84** | **0.060** | **0.17 (0.30)** | **0.15 (0.23)** | **0.10 (2.53)** |
| PHA_TC26 | 17 | 2023 | 25 | 0.419 | 0.742 | 3 | 0.10 | 0.17 | 3 (1–21x) | 0.03 | 0.686 | 0.03 (0.00) | 0.02 (0.00) | 0.27 (2.57) |
| **PHA_SEIsland** | **13** | **2023** | **23** | **0.035** | **0.217** | **8** | **0.33** | **0.70** | **8 (1–12x)** | **0.24** | **0.080** | **0.16 (0.38)** | **0.22 (0.99)** | **-0.33 (9.39)** |
| **PHA_Highway** | **12** | **2023** | **24** | **0.047** | **0.232** | **8** | **0.30** | **0.85** | **8 (1–7x)** | **0.55** | **0.104** | **0.16 (0.44)** | **0.21 (0.82)** | **-0.37 (9.24)** |
| PHA_2W1 | 18 | 2023 | 24 | 0.028 | 0.234 | 15 | 0.64 | 0.95 | 14 (1–4x) | 1.24 | 0.012 | 0.16 (0.31) | 0.16 (0.31) | -0.06 (5.64) |
| **PHA_F12** | **11** | **2023** | **24** | **0.038** | **0.202** | **12** | **0.50** | **0.93** | **12 (1–5x)** | **1.00** | **-0.010** | **0.17 (0.45)** | **0.16 (0.44)** | **0.04 (4.76)** |
| PHA_2NW4 | 19 | 2023 | 24 | 0.041 | 0.225 | 10 | 0.43 | 0.87 | 10 (1–7x) | 0.60 | 0.104 | 0.16 (0.42) | 0.18 (0.55) | -0.14 (6.57) |
| PHA_6NW3 | 20 | 2023 | 24 | 0.120 | 0.438 | 5 | 0.17 | 0.74 | 5 (2–10x) | 0.46 | 0.078 | 0.09 (0.20) | 0.10 (0.31) | -0.10 (7.49) |
| MID_11 | 21 | 2022 | 50 | 0.054 | 0.322 | 17 | 0.33 | 0.86 | 17 (1–13x) | 0.40 | 0.138 | 0.12 (0.20) | 0.10 (0.12) | 0.14 (0.89) |
| MID_13 | 22 | 2022 | 49 | 0.014 | 0.170 | 24 | 0.55 | 0.96 | 23 (1–5x) | 1.14 | 0.047 | 0.19 (0.39) | 0.18 (0.36) | 0.05 (3.63) |
| **MID_33** | **23** | **2022** | **50** | **0.074** | **0.349** | **12** | **0.24** | **0.86** | **12 (1–11x)** | **0.29** | **0.052** | **0.11 (0.24)** | **0.11 (0.27)** | **0.04 (4.83)** |
| MID_87 | 24 | 2022 | 50 | 0.070 | 0.294 | 11 | 0.20 | 0.90 | 11 (1–9x) | 0.60 | 0.014 | 0.13 (0.36) | 0.13 (0.32) | 0.03 (5.69) |
| MID_WP289 | 25 | 2022 | 50 | 0.328 | 0.686 | 8 | 0.15 | 0.43 | 8 (1–36x) | 0.07 | 0.157 | 0.04 (0.03) | 0.03 (0.02) | 0.27 (3.30) |
| MID_R20 | 26 | 2022 | 24 | 0.081 | 0.363 | 10 | 0.41 | 0.84 | 10 (1–8x) | 0.47 | 0.033 | 0.11 (0.21) | 0.07 (0.06) | 0.34 (2.13) |
| **MID_33** | **23** | **2023** | **50** | **0.050** | **0.301** | **18** | **0.35** | **0.91** | **17 (1–11x)** | **0.60** | **-0.001** | **0.13 (0.23)** | **0.13 (0.20)** | **0.00 (3.11)** |
| KUR_R35 | 27 | 2023 | 19 | 0.041 | 0.324 | 8 | 0.41 | 0.80 | 9 (1–7x) | 0.41 | 0.037 | 0.12 (0.22) | 0.13 (0.31) | -0.08 (5.00) |
| KUR_WP013 | 28 | 2023 | 9 | 0.732 | 0.899 | 2 | 0.13 | 0.22 | 2 (1–8x) | 0.06 | 1 | 0.01 (0.01) | 0.01 (0.01) | -0.06 (8.97) |
| KUR_WP018 | 29 | 2023 | 14 | 0.816 | 0.933 | 2 | 0.08 | 0.14 | 2 (1–13x) | 0.03 | 1 | 0.01 (0.00) | 0.01 (0.00) | -0.04 (8.60) |
| **KUR_WP019** | **30** | **2023** | **19** | **0.126** | **0.440** | **7** | **0.35** | **0.78** | **7 (1–8x)** | **0.40** | **0.069** | **0.09 (0.19)** | **0.06 (0.07)** | **0.37 (2.71)** |

### Dataset comparison

We analyzed 1,113 thalli in Dataset One, including reproductive tetrasporophytes at the time of sampling (N = 178), heterozygous multilocus genotypes (MLGs) at one or more loci (N = 808), and/or genotypes that matched a known reproductive tetrasporophyte but was a fixed homozygote (N = 157). We excluded the 47 remaining genotyped thalli from this dataset because they were considered to be gametophytes due to the presence of cystocarps at the time of sampling (n = 8) or were homozygous at all loci and did not match a reproductive tetrasporophytic MLG (n = 39).

In Dataset Two, 866 thalli genotyped were retained because they were reproductive at the time of sampling (N = 178) and/or were heterozygous at one or more loci (N = 808). The 294 remaining thalli that were homozygous fixed and included in Dataset One were considered gametophytes for this dataset.

We found no difference between Dataset One and Dataset Two for ploidy diversity (*P_HD_*, W = 393.5, *p* = 0.680), unbiased probability of identity (*pidu*, W = 531.5, *p* = 0.086), the probability of identity between sibs (*pids*, W = 512, *p* = 0.157), genotypic evenness (*D\**, W = 395.5, *p* = 0.702), the distribution of clonal membership (*pareto ß*, *t_28_ =* -1.85, *p* = 0.075), a multilocus estimate of linkage disequilibrium (*r̄_D_*, W = 406.5, *p* = 0.834), the variance of expected heterozygosity (W = 354, *p* = 0.304), the variance in observed heterozygosity (var[*H_O_*], W = 299, *p* = 0.060), or the variance in the inbreeding coefficient (var[*F_IS_*], W = 316, *p* = 0.106). However, we did find differences between the two datasets in genotypic richness (*R*, *t_28_ =* - 2.755, *p* = 0.010), the multilocus estimates of expected heterozygosity (*H_E_*, *t_28_ =* -2.859, *p* = 0.008), the multilocus estimates of observed heterozygosity (*H_O_*, *t_28_ =* -3.808, *p* = 7.01 × 10^−4^), and the multilocus estimates of the inbreeding coefficient (*F_IS_*, *t_28_ =* 6.683, *p* = 2.97 × 10^−7^).

### Phase ratio

Most *Chondria tumulosa* thalli were vegetative at the time of collection across all sites and time points (∼90%, N = 3,624; Figure 4; Table 1B). Of the thalli that were reproductive (N = 403), 98% were reproductive tetrasporophytes (N = 395) and 2% were cystocarpic female gametophytes (N = 8). Cystocarpic female gametophytes were found at sites on Manawai in July 2021 (N = 4), Kuaihelani in July 2022 (N = 2), and Hōlanikū in July 2023 (N = 2). No reproductive male gametophytes were observed.

**Figure 4.**
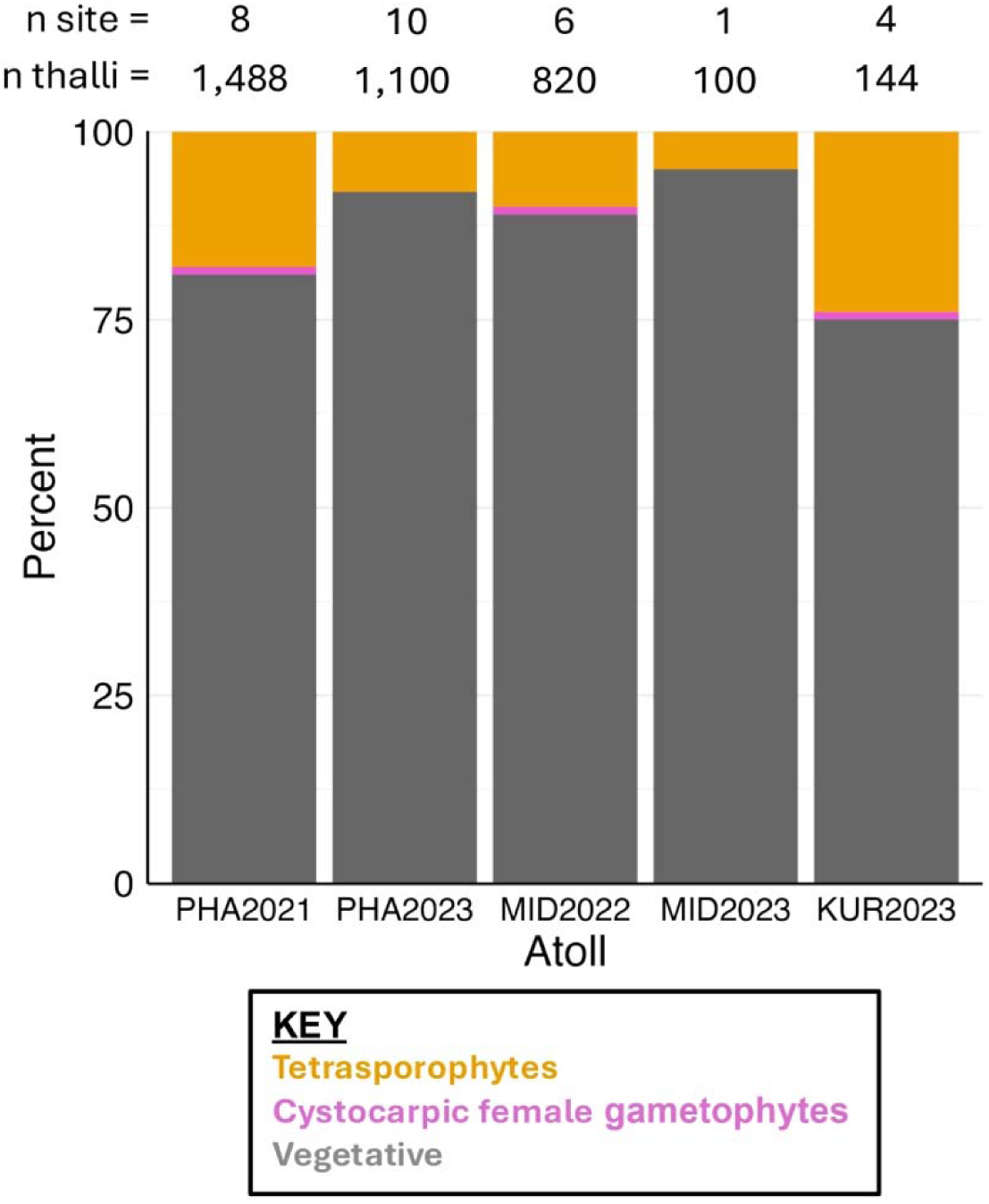
Frequency of *Chondria tumulosa* reproductive tetrasporophytes, cystocarpic female gametophytes, and vegetative thalli across all sampling locations at each atoll by year. No reproductive male gametophytes have been found to date. n site, the number of sites surveyed at each atoll per year; n sample, the number of thalli examined at each atoll per year. Many thalli examined for reproductive state were not later genotyped.

*P_HD_* values were low at most sites and there were significant deviations from the predicted phase ratio of √2:1 (Table 1B, Table S3). For the sites with *P_HD_*values > 0.50, some had small sample sizes (e.g., KUR_WP018 where N = 14) or many fixed homozygous thalli that matched a reproductive tetrasporophyte (e.g., PHA_9.4 where 40 thalli were fixed homozygotes). Observed phase ratios were significantly different than the predicted ratio of √2:1 for all sites at Manawai in 2021 and 2023 (Table 2; Table S3; Table S4A). Only MID_R20 at Kuaihelani was not different from √2:1 in 2022. None of the sites sampled at Hōlanikū differed significantly from the predicted phase ratio, but we note that sample sizes were generally less than 20, likely limiting power to detect small deviations from √2:1 (see Krueger-Hadfield & Hoban, 2016). Unlike all the other sites, KUR_WP013 had more gametophytes based on homozygosity alone where only two thalli were considered tetrasporophytes (Table 1B).

### Genotypic diversity

We found 164 unique genotypes observed out of the 1,113 tetrasporophytes genotyped. Of these 164 unique genotypes, 50 genotypes were re-encountered between 2 and 121 times. Twenty-one of the repeated MLGs had *p_sex_* < 0.05, two had *p_sex_* between 0.05 and 0.06, and the remaining 27 repeated MLGs had *p_sex_* > 0.11 (Table S5). As an example, MLG42 was found 121 times across years and at all atolls (17 sites/time points in total; *p_sex_*= 1.11 × 10^−16^). By contrast, MLG101 was found across Manawai sites in 2021 and 2023 and Kuaihelani MID_33 in 2022 and 2023, but the *p_sex_*was 0.989 (Table S5).

Genotypic richness varied from 0.08 to 0.64 across all sampling sites and time points (Figure 5; Table 2; Figure S1; Table S4B). Genotypic evenness varied from 0.14 to 0.95 with all but four sites being greater than 0.75 (Figure 5, Table 2, Figure S1, Table S4B). There was no clear grouping of sites or time points when comparing the relationship between genotypic evenness (*D\**) and genotypic richness (*R*; Figure 6, Figure S2).

**Figure 5.**
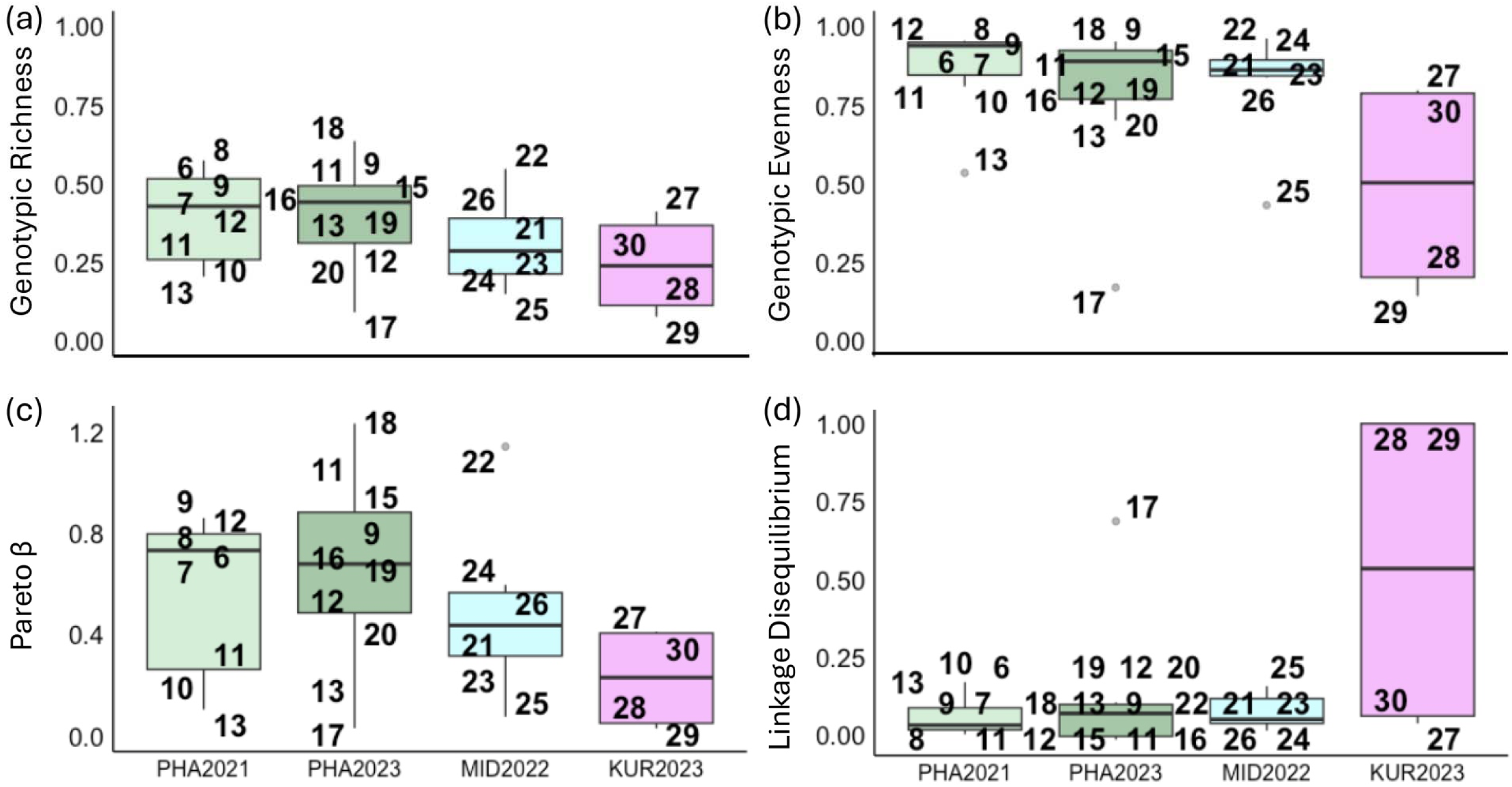
Boxplots by atoll and year for (a) genotypic richness (*R*), (b) genotypic evenness (*D\**), (c) the distribution of clonal membership (*Pareto ß*), and (d) the multilocus estimate of linkage disequilibrium (*r̄_D_*) using Dataset One including all reproductive tetrasporophytes, all heterozygous thalli, and all repeated multilocus genotypes that were identical to a reproductive tetrasporophyte (including fixed homozygous thalli). Boxes represent the interquartile range, the middle lines are medians, whiskers represent the 1.5 interquartile ranges, and the black dots represent outliers. Datapoints are labeled by site number (see Table 1) and jittered along the x-axes to improve visualization.

**Figure 6.**
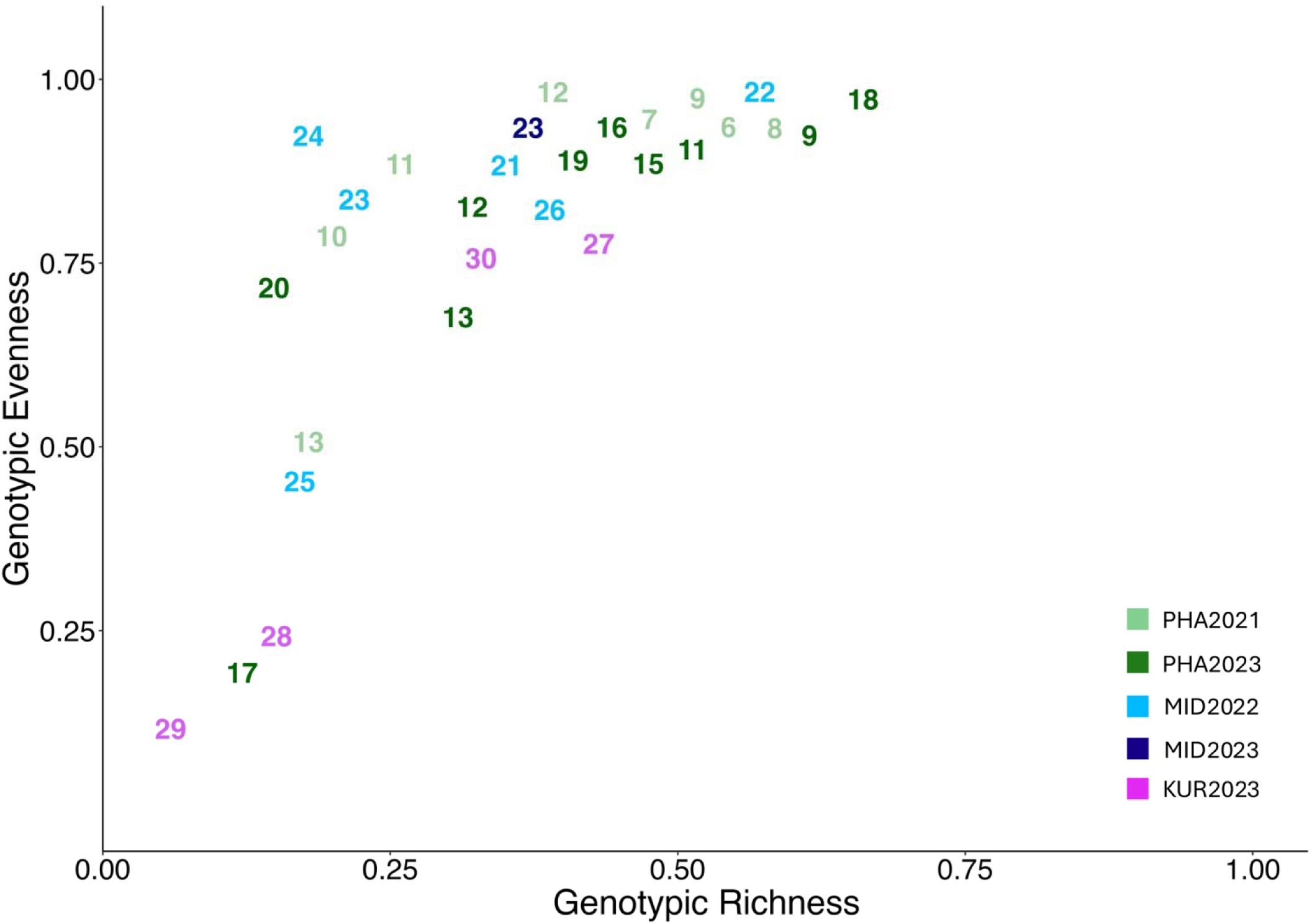
Reproductive mode variation shown as genotypic evenness (*D\**) versus genotypic richness (*R*) based on Dataset One including all reproductive tetrasporophytes, all heterozygous thalli, and all repeated multilocus genotypes that were identical to a reproductive tetrasporophyte (including fixed homozygous thalli). Each number refers to a site in which *Chondria tumulosa* thalli were sampled and are color coordinated by atoll and year (see Table 1).

All sites sampled had *Pareto ß* values < 2. The Manawai site from 2023 PHA_2W1 had the highest *Pareto ß* value of 1.24 (Table 2). More than half the sites and time points had values that were < 0.7 (Table 2, Table S4B).

Linkage disequilibrium (*r̄_D_*) was low for all sites at Manawai and Kuaihelani except for site PHA_TC26 at Manawai in 2023 (Figure 5). At Hōlanikū, the thalli at two sites had smaller *r̄_D_* values (*r̄_D_* < 0.25) while the other two sites sampled had greater *r̄_D_* values (*r̄_D_* > 0.75).

### Genetic diversity

Both expected and observed heterozygosity were low at all atolls across all years (*H_E_* < 0.21; *H_O_* < 0.22; Table 2; Table S4B). The multilocus inbreeding coefficients (*F_IS_*) tended to be positive at most sites with Dataset One (Table 2), with many values at 1 (Figure 7). However, most sites in Dataset Two were characterized by negative multilocus *F_IS_* values (Table S4B). The variance in *F_IS_* across loci was high at all sites and across all time points. Manawai sites in 2021 exhibited the greatest variance and Kuaihelani sites in 2022 had the smallest variance (Figure 7; Figure S3; Table S6A, Table S6B).

**Figure 7.**
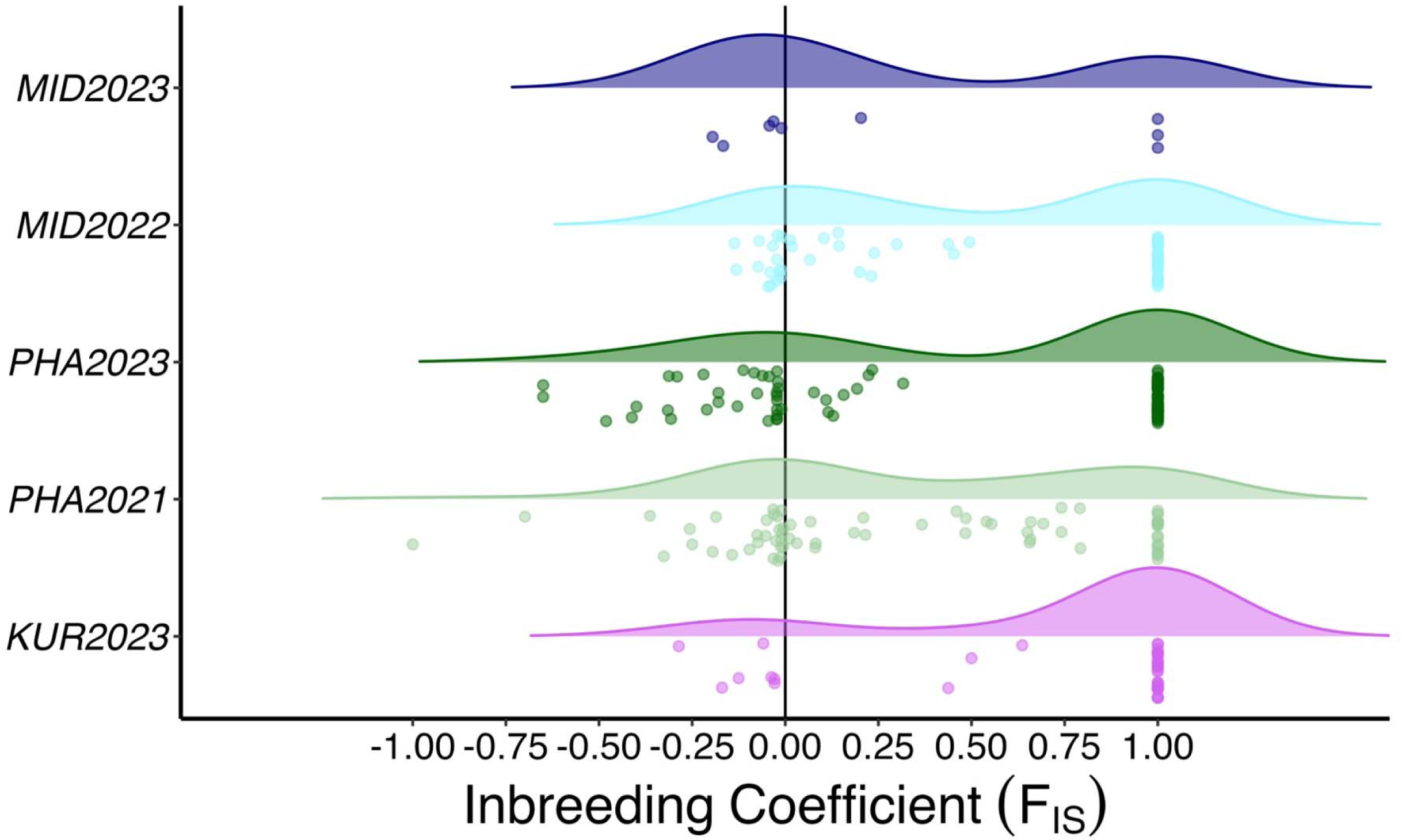
Density plot of single locus estimates of the inbreeding coefficient (*F_IS_*) by atoll and year for sites in which we sampled *Chondria tumulosa* thalli. *F_IS_* was calculated on Dataset One including all reproductive tetrasporophytes, all heterozygous thalli, and all repeated multilocus genotypes that were identical to a reproductive tetrasporophyte (including fixed homozygous thalli). Number of data points representing single locus *F_IS_* values for each site, atoll, and year are as follows: MID2023 (9), MID2022 (54), PHA2023 (90), and PHA2021 (72), KUR2023 (36).

### Temporal analyses of reproductive mode

Changes in the resulting genotype frequencies inferred relative rates of outcrossing, selfing, and clonality which were found to be variable depending on the site (Figure 8, Figure 9; Table 3; Table S7). The only site that was sampled at Kuaihelani at more than one timepoint was MID_33, and its inferred clonal rate was zero. The rates of outcrossing and selfing remained somewhat consistent, though the site had an outcrossing rate of 1 between July 2022 and 2023 (Figure 8). We observed much greater variation in the reproductive mode as assessed by temporal patterns of genotype frequencies at the four sites sampled in July 2021 and July 2023 on Manawai. PHA_F12 and PHA_F4 were characterized by high outcrossing rates (Figure 9). In contrast, the PHA_SE Island site had selfing and clonal rates each equal to 0.5. Finally, PHA_Highway had a clonal rate equal to 1.

**Figure 8.**
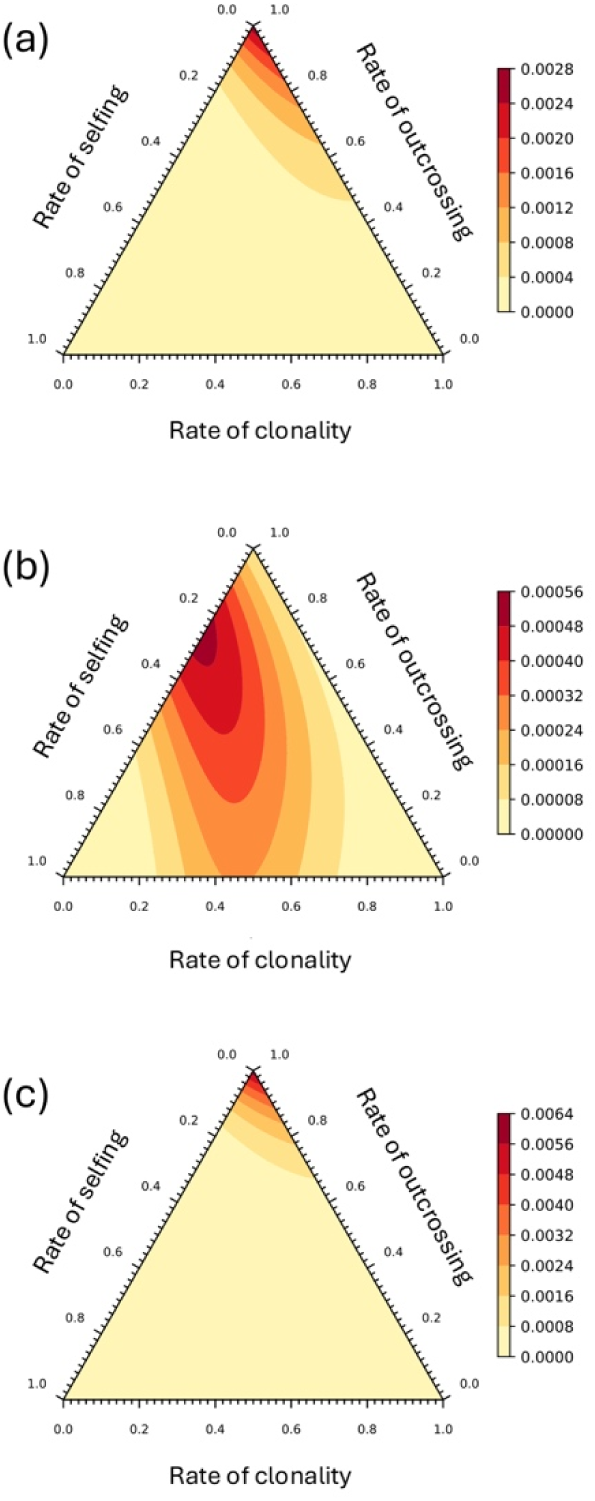
The rates of selfing, outcrossing, and clonality for the site MID_33 (#23, Kuaihelani, Table 1) at which we sampled *Chondria tumulosa* thalli (a) in July 2022 and July 2023, (b) in July 2022 and October 2022, and (c) in October 2022 and July 2023 (see also Table 4). We used Dataset One, including all reproductive tetrasporophytes, all heterozygous thalli, and all repeated multilocus genotypes that were identical to a reproductive tetrasporophyte (including fixed homozygous thalli).

**Figure 9.**
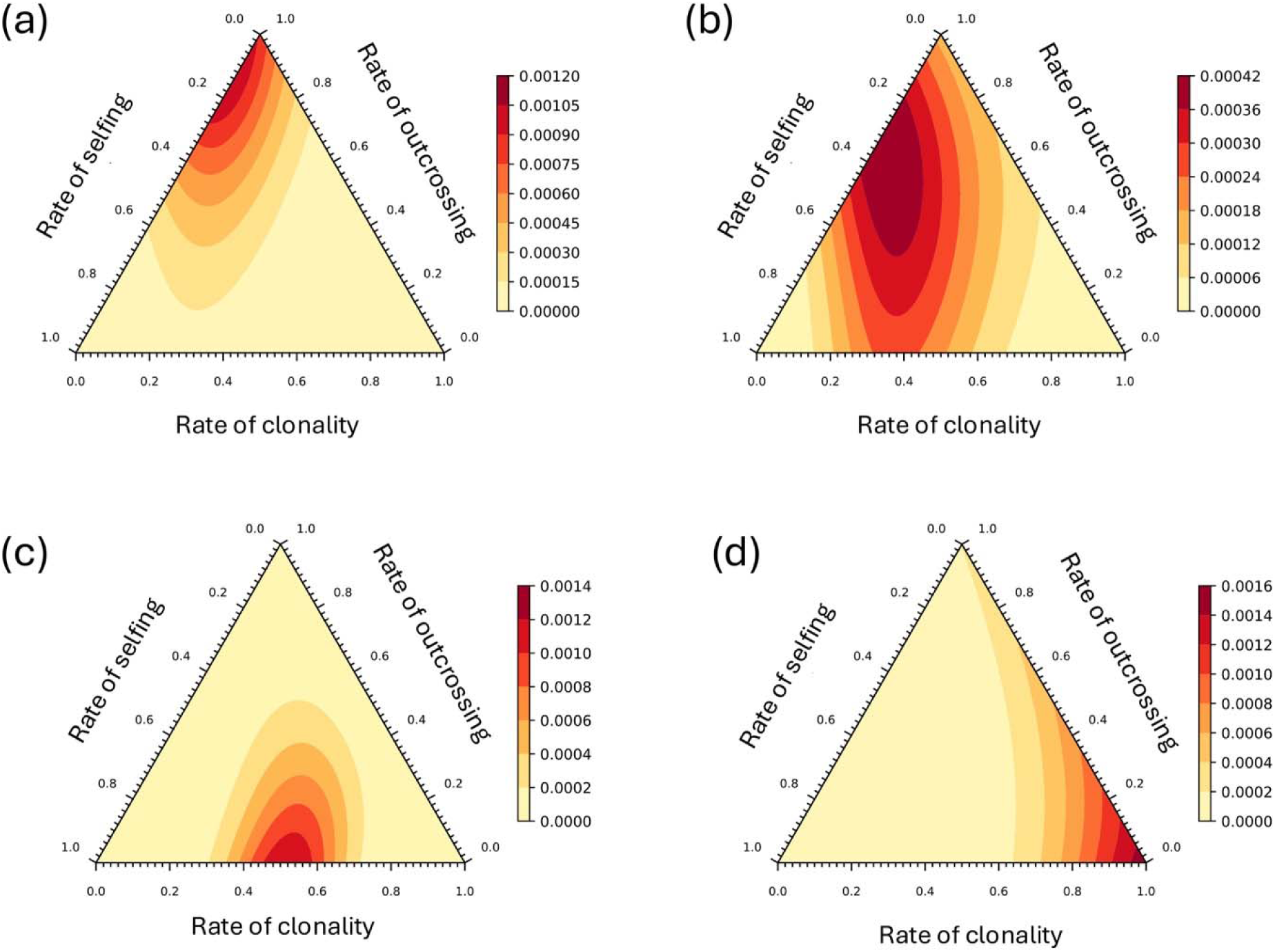
The rates of selfing, outcrossing, and clonality for sites at Manawai for which we sampled *Chondria tumulosa* thalli in July 2021 and July 2023. (a) Manawai site PHA_F12 (#11; Table 1), (b) Manawai site PHA_F4 (#9), (c) Manawai site PHA_SEIsland (#13), and (d) Manawai site PHA_Highway (#12; see also Table 4). We used Dataset One, including including all reproductive tetrasporophytes and all heterozygous thalli.

**Table 3.** Credible intervals of inferred rates of outcrossing, selfing, and clonality from the genotype frequencies at sites in which we sampled *Chondria tumulosa* thalli in at least two time points. We used Dataset One, including all reproductive tetrasporophytes, all heterozygous thalli, and all repeated multilocus genotypes that were identical to a reproductive tetrasporophyte (including fixed homozygous thalli).

| Site | Site # | Comparison | Outcrossing | Selfing | Clonality |
| --- | --- | --- | --- | --- | --- |
| MID_33 | 23 | July 2022 to October 2022 | 0.74 (0,1) | 0.26 (0,0.81) | 0.00 (0,0.81) |
| MID_33 | 23 | July 2022 to July 2023 | 1.00 (0,1) | 0.00 (0,0.47) | 0.00 (0,1) |
| MID_33 | 23 | October 2022 to July 2023 | 1.00 (0.13,1) | 0.00 (0,0.38) | 0.00 (0,0.8) |
| PHA_F4 | 9 | July 2021 to July 2023 | 0.64 (0,1) | 0.33 (0,0.87) | 0.03 (0,0.74) |
| PHA_F12 | 11 | July 2021 to July 2023 | 0.86 (0,1) | 0.14 (0,0.85) | 0.00 (0,0.49) |
| PHA_Highway | 12 | July 2021 to July 2023 | 0.00 (0,1) | 0.00 (0,0.51) | 1.00 (0,1) |
| PHA_SEIsland | 13 | July 2021 to July 2023 | 0.00 (0,0.8) | 0.48 (0.01,0.8) | 0.52 (0.03,0.8) |

### Ranking sites by clonality and selfing

Based on a combination of summary statistics and ranking, all sites exhibited signatures suggesting very highly clonal populations (Table 2, Table S8). The normalized and categorical ranks were highly correlated (Figure S8), adding confidence in the ranking of the sites. Using Σ.cln, the sums of the normalized population genetic values accounting for importance of clonality, sites at Hōlanikū tended to be more clonal than Manawai and Kuaihelani (H=3.83, p=0.1473) but not significantly (Hōlanikū - Manawai: p = 0.167 and Hōlanikū - Kuaihelani: p = 0.199) while Manawai and Kuaihelani were similar in their clonal ranks (p = 0.888).

### Genetic structure

Within and among atolls, we found evidence of strong genetic structure in *Chondria tumulosa* (Table 4A, Table S9). Manawai had a mean *F_ST_*value of 0.186 with values ranging from 0.001 ≤ *F_ST_* ≤ 0.632. Kuaihelani had a mean *F_ST_* value of 0.112 with values ranging from 0.000 ≤ *F_ST_* ≤ 0.372. Hōlanikū had the highest genetic differentiation with a mean *F_ST_* value of 0.536 and values ranging from 0.122 ≤ *F_ST_* ≤ 0.954. Genetic differentiation between atolls in 2023 was greater than genetic differentiation within atolls (Table 4A). The 2021 Manawai and 2022 Kuaihelani sampling events had comparable levels of within-atoll differentiation. Mantel tests using both direct and indirect geographic distance calculations revealed increased genetic differentiation with increasing geographic distance (Figure S5, Table S10).

**Table 4A.** Mean *F_ST_* values from pairwise comparisons between sites within an atoll and across atolls in the same collection year in 2023 where we collected *Chondria tumulosa* thalli. We used Dataset One, including all reproductive tetrasporophytes, all heterozygous thalli, and all repeated multilocus genotypes that were identical to a reproductive tetrasporophyte (including fixed homozygous thalli).

| Location | Number of Pairwise Comparisons | Mean $F_{ST} \pm SE$ | Range |
| --- | --- | --- | --- |
| PHA2021 | 28 | $0.181 \pm 0.025$ | 0.010 – 0.464 |
| PHA2023 | 45 | $0.215 \pm 0.025$ | 0.010 – 0.632 |
| PHA All | 153 | $0.186 \pm 0.012$ | 0.001 – 0.632 |
| MID2022 | 15 | $0.149 \pm 0.029$ | 0.008 – 0.372 |
| MID All | 28 | $0.112 \pm 0.021$ | 0.000 – 0.372 |
| KUR2023 | 6 | $0.513 \pm 0.102$ | 0.122 – 0.954 |
| KUR2023–MID2023 | 4 | $0.279 \pm 0.133$ | 0.072 – 0.594 |
| KUR2023–PHA2023 | 40 | $0.368 \pm 0.038$ | 0.003 – 0.803 |
| PHA2023–MID2023 | 10 | $0.197 \pm 0.047$ | 0.018 – 0.478 |

Strong genetic structure was evident through time both within and between atolls (Figure 10; Figure S6; Figure S7). The global stepping-stone shape of the minimum spanning tree of pairwise genetic distances indicates that relatedness between individuals was hierarchically structured by atolls and then by years, following global geographic distances. The thalli collected from Manawai (PHA) in 2019, 2021, and 2023 clustered closely together on the minimum spanning tree, but each year was distinct (Figure 10). Genetic differentiation between years (2021 and 2023) at sites that were sampled at both time points was low (PHA_F4, PHA_F12, and PHA_Highway) and remained stable along years between atolls (Figure S7; Table 4B). Distributions of genetic differentiation remained stable from 2021 to 2023 in all sites (PHA_F4 2021 versus 2023: U=29, p=0.62; PHA_F12 2021 versus 2023: U=35, p=0.38; PHA_Highway 2021 versus 2023: U=29.5, p=0.60), except for PHA_SEIsland (2021 versus 2023: U=49, p=0.04) in which genetic differentiation significantly decreased from 2021 to 2023 (Table S11).

**Figure 10.**
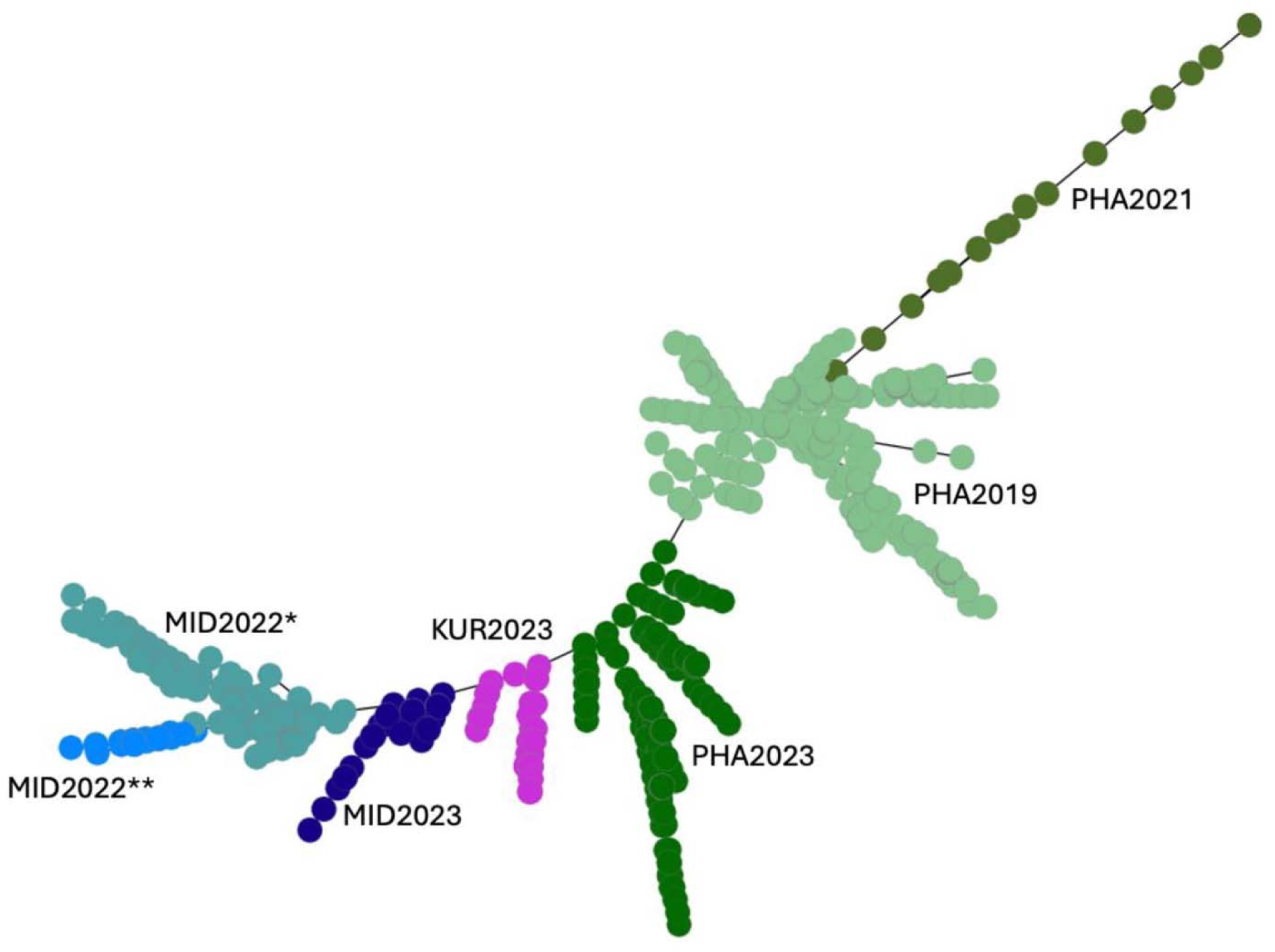
Minimum spanning tree for the sites in which we sampled *Chondria tumulosa* thalli using Dataset One, including all reproductive tetrasporophytes, all heterozygous thalli, and all repeated multilocus genotypes that were identical to a reproductive tetrasporophyte (including fixed homozygous thalli). *Sampled in July 2022; **Sampled in October 2022

**Table 4B.**
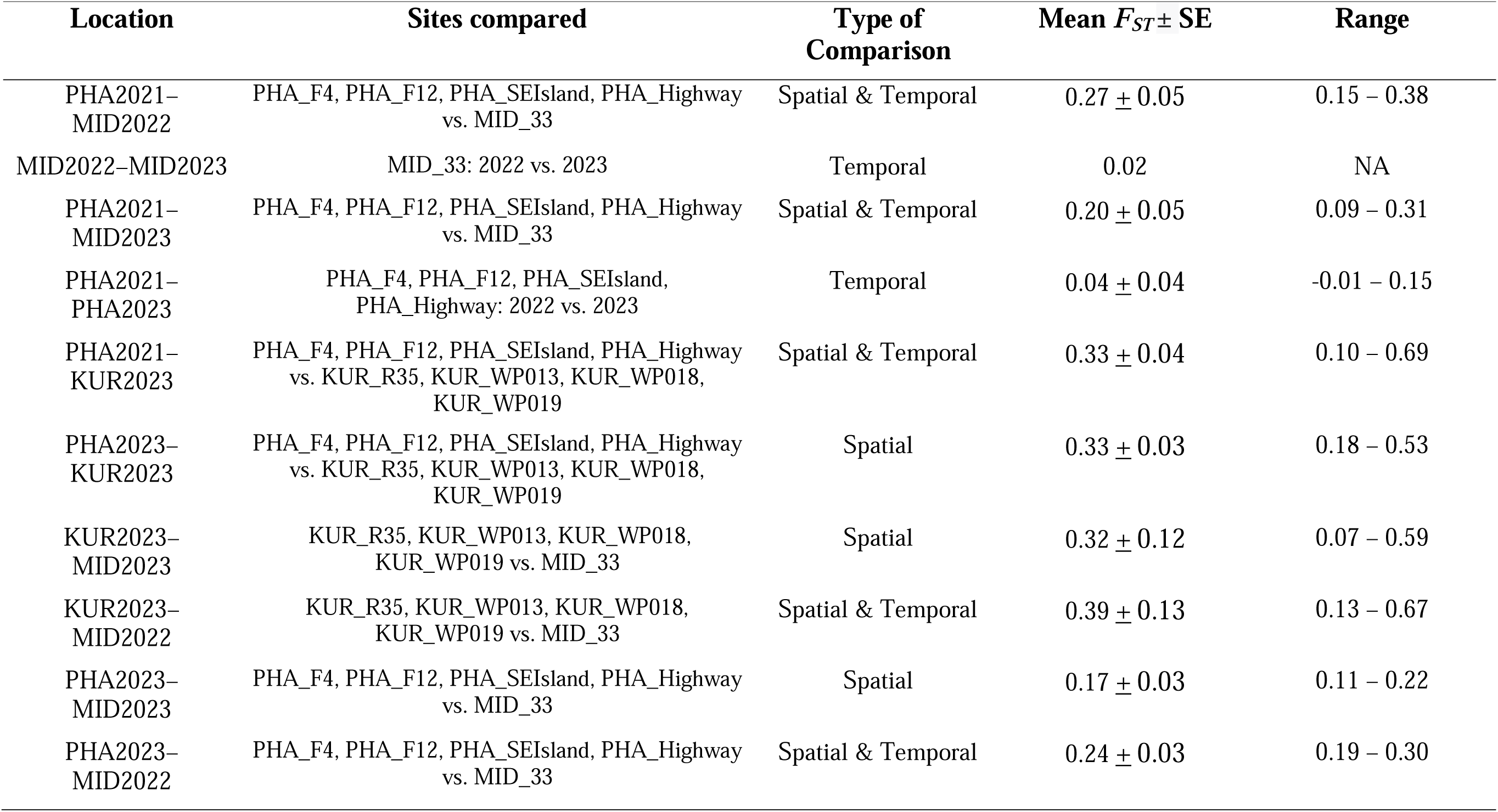
Mean *F_ST_* values from pairwise comparisons within an atoll and across atolls using sites that *Chondria tumulosa* thalli were collected at multiple time points for the temporal analyses (Manawai, Kuaihelani (MID_33)) and at a single time point for the spatial analyses (Manawai, Kuaihelani, and Hōlanikū). We used Dataset One, including all reproductive tetrasporophytes, all heterozygous thalli, and all repeated multilocus genotypes that were identical to a reproductive tetrasporophyte (including fixed homozygous thalli).

## DISCUSSION

In this study, we expanded on our previous research focused on a single sampling event at several sites at Manawai (PHA) in 2019 (Williams et al., 2024) by including additional sites at multiple atolls across several years, covering the known distribution of this alga to date. This intensive spatial and temporal sampling provides a more thorough understanding of the genetic structure across and within the atolls. Tetrasporophytes dominated at all sites, atolls, and throughout the sampling period from 2019 to 2023 – a hallmark of some free-living algae (see descriptions of algal population types in Krueger-Hadfield et al., 2023) and recent colonization and expansion dynamics (see Krueger-Hadfield, 2020). We did, however, find eight cystocarpic female gametophytes at the time of collection, and these females were haploid based on their microsatellite multilocus genotypes. While reproductive male gametophytes have not yet been observed, the presence of cystocarps suggests that males may have been present at times when we were not able to sample. While we have limited statistical power with our current suite of microsatellite loci (see *pidu* and *pids* values as well as *p_sex_*values), we consider each site to be highly clonal. The relative values of the summary statistics coupled with low *pid* could also suggest high rates of intergametophytic selfing (Klekowski, 1969; see Krueger-Hadfield et al., 2024; Krueger-Hadfield, 2024 for more discussion on selfing in dioicous gametophytes) when sexual reproduction does occur or very low local effective population sizes. Moreover, the haploid proportion is likely very small, with a concomitant strong influence on genetic diversity (see Stoeckel et al. 2021). The effects of clonality, intergametophytic selfing, and small haploid proportions on genetic diversity are likely further exacerbated by huge biomass fluctuations (see Lopes et al., 2023) that lead to extinction and recolonization dynamics, resulting in strong genetic drift. As a result, our single time point genotyping provided better resolution of variation in the reproductive mode than the temporal genotyping. Unsurprisingly, we found strong genetic structure across sites sampled at Manawai, Kuaihelani, and Hōlanikū. While the directionality of spread for *Chondria tumulosa* is uncertain, we found strong isolation by distance which matched geographical distances between atolls. In the following sections, we discuss these results and their consequences for the management of *Chondria tumulosa* in PMNM, and more broadly for other haploid-diploid macroalgae.

### Comparison between datasets

All patterns we observed were interpreted with caution due to the rather poor *pid* values, thereby limiting our ability to robustly distinguish between genotypes. This issue likely stems from insufficient polymorphism in the current suite of loci, itself reflecting limited genetic diversity and ancestries at the time of the initial colonization. As a result, repeated MLGs were observed across both time and space, despite the pairwise genetic distances and *F_ST_* values suggesting strong genetic structure. Indeed, more than half of the re-encountered MLGs had *p_sex_*much greater than the nominal cut-off of 0.05. At some sites, such as the Manawai sites sampled in both 2021 and 2023 (e.g., PHA_F4), we might expect to re-encounter an MLG if the MLG in question persists at a site (see as an example the temporal study of genotypic diversity in the red alga *Gracilaria vermiculophylla*, Oetterer et al., 2025). However, at present, we cannot distinguish whether these repeated MLGs are the same or a product of low statistical power.

Increasing the number of loci would not only increase statistical power but could also help resolve between phases in the life cycle. We observed an increase in homozygosity from our original sampling in 2019 (Williams et al., 2024) to our current study across several loci. Without an alternative and complementary method to distinguish between phases and sexes, such as sex-linked markers (e.g., Krueger-Hadfield et al., 2021b), we might have inadvertently included some gametophytes in Dataset One due to our limited power. Nevertheless, most of our summary statistics were the same between the two datasets. Importantly, ploidy diversity was not different between the two datasets, suggesting our observations of tetrasporophytic bias were robust. The significant differences between the two datasets made intuitive sense. Genotypic richness, expected heterozygosity, and observed heterozygosity increased with Dataset Two because this data set did not have the large number of repeated and homozygous thalli included. Similarly, the inbreeding coefficient became more negative across sites in Dataset Two, as thalli were generally more heterozygous. While future studies should use more loci, such as single nucleotide polymorphisms (e.g., see methods of Delord et al., 2018 implemented in Besnard et al., 2023), to improve statistical power for the resolution between phases and genotypes, the patterns we discuss here are generally robust based on all results.

### Prevailing reproductive mode

At the time of sampling, most *Chondria tumulosa* thalli were vegetative. When non-vegetative thalli were found, they were overwhelmingly tetrasporophytes (diploids), apart from the eight cystocarpic female thalli (< 1%). Other studies have found tetrasporophytic bias was due to artificial selection from algal cultivation (Guillemin et al., 2008), selective pressures of the invasion of soft sediment habitats (Krueger-Hadfield et al., 2016), or changes in salinity (Gabrielson et al., 2002). Thus, we may posit that there are ecological conditions that select for the tetrasporophytic phase of the life cycle after which clonal processes through thallus fragmentation led to the maintenance of a tetrasporophytic bias. In *Gracilaria chilensis*, vegetative tetrasporophytes grew fastest, providing an explanation as to why farmers have artificially selected for this phase (Guillemin et al., 2013). Diploidy may buffer against deleterious mutations in small pioneering populations enduring repeated foundation events, may help maintain more alleles that would enhance adaptive potential in new environments by recombination, and may provide higher fitness under stressful conditions commonly found in angiosperms (Otto & Goldstein, 1992; Jenkins & Kirkpatrick, 1995; Madlung 2013; Turcotte et al. 2024). Additionally, diploidy and vegetative reproduction facilitate the maintenance of heterozygosity, which can increase fitness through mechanisms like heterosis (Stoeckel et al., 2021). If similar differences in growth rates occur between *C. tumulosa* gametophytes and tetrasporophytes, this could explain the patterns we observed, though this remains to be tested.

Previous macroalgal studies have demonstrated the usefulness of temporal sampling using the comparison of genotype frequencies across two time points (Becheler et al., 2017) in understanding the relative contributions of sexual and clonal reproduction (Heiser et al., 2023b; Thornton et al., 2024). For example, *Avrainvillea lacerata* blades collected from O ahu between 2018 and 2023 were used for temporal comparisons of genotype frequencies and resulted in the detection of high clonal rates (Thornton et al., 2024). This alga spreads largely through vegetative reproduction (Thornton et al., 2024), corroborating earlier ecological observations (Smith et al., 2002; Veazey et al., 2019). In *Plocamium* spp. from Antarctica (see also Heiser et al., 2025), Heiser et al. (2023b) found variation at two sites in which one site had a high rate of selfing while the other had a high clonal rate with background selfing (Heiser et al., 2023b). The summary statistics calculated in each of these sites matched the selfing and clonal rates observed between single time point and temporal sampling. However, the temporal patterns for *Chondria tumulosa* did not align with prior findings for this species (Williams et al., 2024) or with the patterns observed across the summary from single time point estimates (this study). High clonal rates were expected at the sites sampled over two periods at Manawai and Kuaihelani. Yet, we observed variation in clonality, selfing, and outcrossing within a single atoll – Manawai – that we would expect to see at an interspecific level. We do not know the generation time for *C. tumulosa* and it is therefore likely that our sampling events separated by two years violated some of the assumptions of this method. At Kuaihelani, however, the thalli were collected in July 2022, October 2022, and July 2023, a timeframe likely much closer to representing two consecutive generations. The changes in genotype frequencies were suggestive of complete outcrossing, contradicting all singe time point estimates. This may be due to wide local variations of genetic diversity putatively caused by strong genetic drift or extinction-recolonization dynamics. Our inability to use these temporal genotypic comparisons may be further exacerbated by biomass fluctuations observed at *C. tumulosa* sites when seasonal wave motion removes large mats of interwoven thalli (Lopes et al., 2023). Such fluctuations may generate regular extinction and recolonization events, in which genetic diversity fluctuates due to genetic drift (Nei et al., 1975) coupled with a very small haploid proportion (see discussion in Stoeckel et al., 2021).

Based on this combination of observations of reproductive structures, population genetic summary statistics, and ranking sites by clonality, all sites in which *Chondria tumulosa* has been sampled thus far are highly clonal. The site with the lowest ranking (MID_R20 at Kuaihelani) was a site characterized by shallow depth (∼3 m) and notably very low percent cover of *C. tumulosa* (∼5%). Conversely, PHA_SEIsland and PHA_Highway, both of which are characterized by drifting thalli and high percent cover (70% and 100%, respectively), tended to be among the sites with the highest clonal ranking. Comparable mat-forming macroalgae with high biomass, such as *Eucheuma/Kappaphycus* spp. in Kane’ohe Bay on O’ahu (Woo, 2000; Smith et al., 2002), have been suggested to reproduce clonally in a similar manner to *C. tumulosa* at PHA_SEIsland and PHA_Highway. However, studies using population genetic tools with *Eucheuma/Kappaphycus* spp. thalli are needed to confirm these observations.

### Genetic structure and origins of Chondria tumulosa in PMNM

The native range of *Chondria tumulosa* remains unknown (Sherwood et al., 2020). Our observations to date, such as tetrasporophytic bias, are strongly suggestive of a recent introduction (Williams & Smith, 2007; Krueger-Hadfield, 2020). However, at Hōlanikū, we questioned whether *C. tumulosa* could be an endemic macroalga in low abundance with the potential to exhibit bloom dynamics, such as *Hypnea cervicornis* at other atolls within PMNM (T.M. Williams & B.B. Hauk, *personal observations*). Unlike the large accumulations of biomass observed at Kuaihelani and Manawai, *C. tumulosa* thalli at Hōlanikū appear to integrate into the local algal assemblage in a manner more consistent with native macroalgae, such as *Laurencia* sp. or *Chondrophycus* sp. (T.M. Williams, B.B. Hauk, & H.L. Spalding, *personal observations*). Moreover, Hōlanikū is an atoll known for its high number of endemic species (Kane et al., 2014; Sherwood & Guiry, 2023). Thus, our observations at this atoll raised the possibility that *C. tumulosa* existed in low abundance throughout the PMNM before environmental shifts, such as increased upwelling (A. Kealoha, *personal communication*), triggered its expansion at Kuaihelani and Manawai. A similar phenomenon was observed in 2008 when several consecutive weeks of low wind and warm sea surface temperatures led to *Boodlea composita* blooms, a normally non-blooming macroalga, in the shallow water lagoons at Kuaihelani and Hōlanikū (Vroom et al., 2009). However, in our study, population genetic signatures at Hōlanikū are strongly suggestive of clonal processes, including low *Pareto ß* values, and a more recent introduction. Future work will need to investigate why the ecology of *C. tumulosa* is different at Hōlanikū as compared to the two other atolls in the PMNM.

The first report of *Chondria tumulosa* was in 2016 (Sherwood et al., 2020), coinciding potentially with the arrival of marine debris from the 2011 Tōhoku earthquake and subsequent tsunami (Murray et al., 2018). The first debris reached O ahu in September 2012 (DLNR DAR, *unpublished data*) and has been reported washing up in other areas of the Pacific (Hansen et al., 2019). Observations of Japanese marine debris attributed to this event have been reported in Oregon and Washington state between June 2012 to July 2016, many of which carried fouling marine algae originating from Japan– 83% of which were reproductive at the time (Hansen et al., 2019). Even earlier, the 2004 tsunami in Indonesia likely produced comparable, if not greater debris; however, this event was poorly documented as are the downstream impacts. We have observed *C. tumulosa* thalli re-attached to derelict fishing nets at Manawai (T.M. Williams, B.B. Hauk, & H.L. Spalding, *personal observations*). Manawai is the largest atoll in PMNM with a large inner lagoon containing a structurally complex coral reef system which creates a known marine debris accumulation area. From introduction at Manawai, *C. tumulosa* thalli likely spread to Kuaihelani and Hōlanikū, facilitated by marine debris as well as its association with other algae known to raft. For instance, we frequently observed *C. tumulosa* thalli growing epiphytically on the holdfast and stipes of the brown alga *Turbinaria ornata* (Figure 2). *Turbinaria ornata* thalli have been observed floating both within and between atolls due to the buoyant nature of their pneumatocysts. If *Turbinaria ornata* thalli are a possible vector for *C. tumulosa* thalli, particularly for long-distance transport, then this might help explain how small, negatively buoyant *C. tumulosa* thalli and short-lived spores are able to spread across the northernmost atolls in the PMNM. Nevertheless, strong isolation by distance was observed and match patterns observed using comparable molecular markers in other red algae (e.g., Engel et al., 2004; Krueger-Hadfield et al., 2017; see also Durrant et al., 2014).

*Chondria tumulosa* sites exhibited both temporal and spatial differentiation. For example, at Manawai, genetic differentiation remained stable from 2021 to 2023 except in SEIsland. The loss of strong genetic structure over time is expected for introduced species where founder effects are responsible for initial differentiation (Dlugosch & Parker, 2008). While clonal reproduction would maintain this differentiation, intergametophytic selfing, when and if sexual reproduction occurs, would erode diversity (Charlesworth & Wright, 2001).

### Future directions for management and conservation

The PMNM is one of the most remote island archipelagos in the world and is managed by four co-trustees: the National Oceanic and Atmospheric Administration, the U.S. Fish and Wildlife Service, the Department of Land and Natural Resources, and the Office of Hawaiian Affairs. The remote nature and complex management structure of the PMNM present unique challenges for conservation and invasive species management. Algal blooms of native algae have occurred in PMNM, including *Boodlea composita* (Vroom et al., 2009) and *Hypnea* sp. (Tsuda et al., 2015), these species are native to the region and were not considered to be invasive species at the time. In contrast, the red alga *Chondria tumulosa* is unique in PMNM with no previous record of occurrence. Since its discovery, it has rapidly established itself and become the dominant macroalga in the northernmost atolls. Progress has been made in understanding its biology, including its distribution (Lopes et al., 2023; Opunui, 2023), ecology (Kaluhiokalani, 2024), and reproductive mode (Williams et al., 2024; this study). Understanding the relative contributions of sexual versus clonal reproduction is crucial for developing effective management strategies to prevent spread between atolls and for anticipating its potential to evolve in the next decade. *Chondria tumulosa* can spread through debris, derelict gear, and other algae, subsequently re-attaching to the substrate by getting caught in the vertical relief of the reef or by epiphytizing other macroalgae. Coupled with its capacity to fragment, there is potential for spread to other atolls as well as the Main Hawaiian Islands. Preventing *C. tumulosa* thalli from crossing the biogeographic break separating these northwestern atolls from the remainder of the Hawaiian Archipelago is critical for protecting coral reef systems across the region (see biogeographic patterns described in Toonen et al., 2011).

While this study has provided valuable insights into the genetic structure of *Chondria tumulosa* throughout its distribution in PMNM, future research efforts should focus on hierarchical sampling strategies as well as increasing the number of loci across the genome to refine patterns of gene flow. Spatial sampling conducted for numerous kelp species have found that small-scale processes were important to structuring populations (Alberto et al., 2010; Billot et al., 2003; Coleman et al., 2011). Importantly, documenting the tropical red algal flora in the Pacific is critical both for taxonomic inventories and to determine the origin of *C. tumulosa* thalli, as has been done to determine the origin of *Gracilaria vermiculophylla* (Krueger-Hadfield et al., 2017) and its environmental tolerances (Sotka et al., 2018). Continued monitoring is critical to document the population dynamics of *C. tumulosa* at Manawai, Kuaihelani, and Hōlanikū, as well as other atolls that could be inundated with thalli in the future.

## ACKNOWLEDGEMENTS

We thank M. Crowley and A. Cao at the Heflin Center for Genomic Sciences for performing fragment analysis; K. Lopes, J. Leonard, K. Peyton, and K. Rankin for help with sample collection; and the NOAA team and affiliates at Papahānaumokuākea National Marine Sanctuary and the USFWS team at Midway National Wildlife Refuge for help with field logistics and resource access. This project was supported by an International Phycological Society Paul Silva Student Grant (to TMW), start-up funding from the College of Arts and Science at the University of Alabama at Birmingham (UAB; to SAKH), the National Fish and Wildlife Foundation (#0810.20.068602 to HLS), the National Fish and Wildlife Foundation Papahānaumokuākea Research and Conservation Fund (#0810.20.068622 and #0810.20.074235 to HLS and SAKH), Clonix2D ANR-18-CE32-0001 (to SS and SAKH), and the Professional Development Exchange from the NSF Research Coordinated Network for Evolution in Changing Seas (OCE-1764316). TMW was supported by the Nancy Foster Scholarship provided by NOAA’s National Marine Sanctuaries. SAKH was supported by an NSF CAREER Award (DEB-2141971 [UAB] and DEB-2436117 [VIMS|WM]). All scientific collections were made under permit numbers PMNM-2021-019 and PMNM-2022-011.

## AUTHOR CONTRIBUTIONS

**Taylor M. Williams** Conceptualization (supporting), Data curation (equal), Formal Analysis (equal), Funding acquisition (supporting), Investigation (equal), Visualization (equal), Writing – original draft (equal), Writing – review and editing (equal); **Heather L. Spalding** Conceptualization (equal), Data curation (supporting), Funding acquisition (equal), Investigation (equal), Resources (equal), Supervision (supporting), Visualization (supporting), Writing – original draft (supporting), Writing – review and editing (equal); **Solenn Stoeckel** Formal analysis (equal), Funding acquisition (supporting), Investigation (supporting), Methodology (equal), Software (lead), Writing – original draft (equal), Writing – review and editing (equal); **Brian B. Hauk** Investigation (supporting), Writing – review and editing (equal); **Jonathan H. Plissner** Investigation (supporting), Writing – review and editing (equal); **Randall K. Kosaki** Investigation (supporting), Writing – review and editing (equal); **Stacy A. Krueger-Hadfield** Conceptualization (equal), Data curation (equal), Formal Analysis (equal), Funding acquisition (equal), Investigation (equal), Methodology (equal), Project Administration (lead), Resources (equal), Software (supporting), Supervision (lead), Visualization (equal), Writing – original draft (equal), Writing – review and editing (equal)

## ABBREVIATIONS

PMNM: Papahānaumokuākea Marine National Monument
MLG: Multilocus Genotype
PCR: Polymerase Chain Reaction
PHA: Pearl and Hermes Atoll (Manawai)
MID: Midway Atoll (Kuaihelani)
KUR: Kure Atoll (Hōlanikū)
UNESCO: United Nations Educational, Scientific, and Cultural Organization
BSA: Bovine Serum Albumin
NOAA: National Oceanic and Atmospheric Administration
FWS: U.S. Fish and Wildlife Service
DLNR: Department of Land and Natural Resources

## SUPPLEMENTARY FIGURES

**Figure S1.**
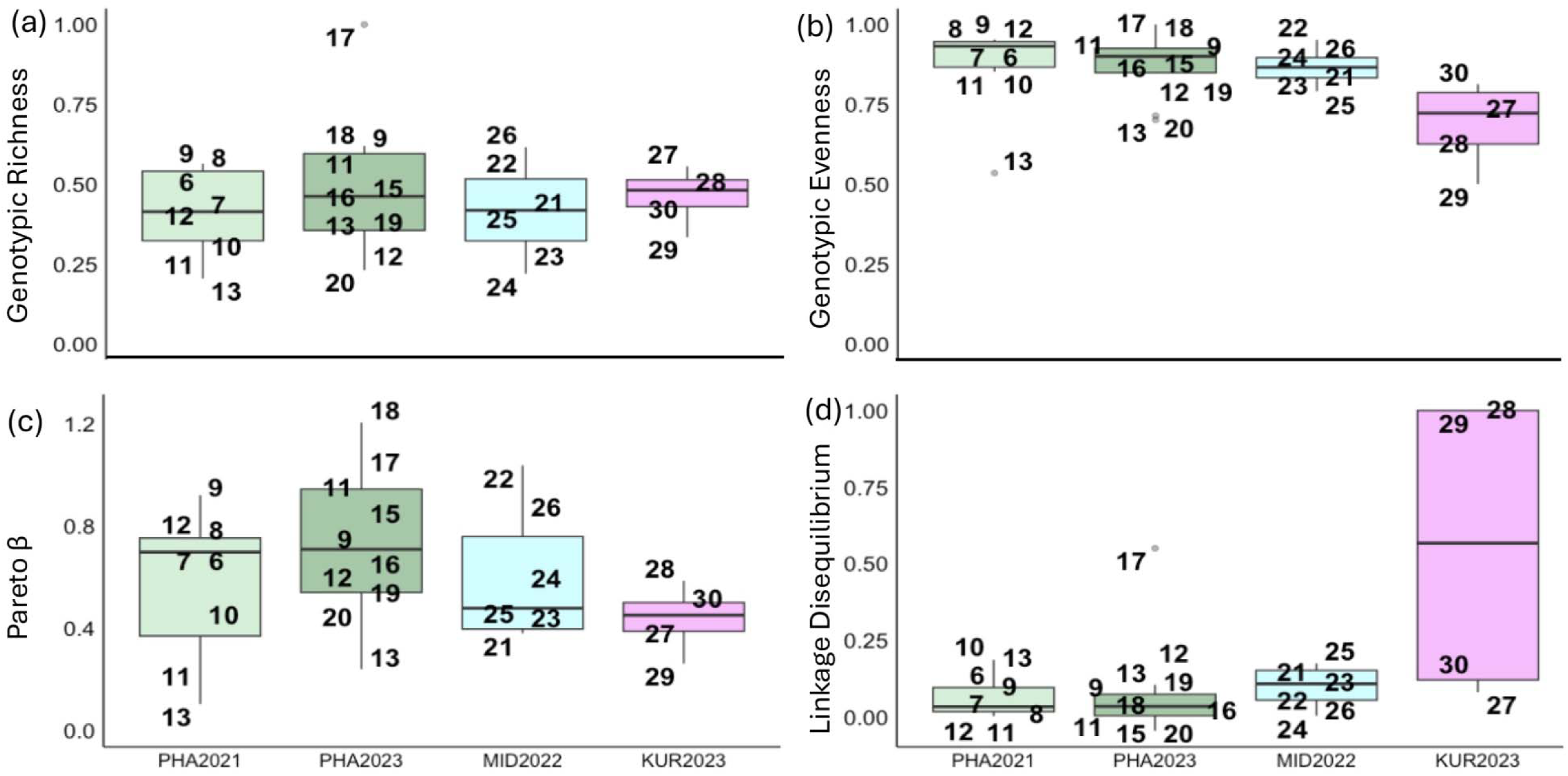
Boxplots by atoll and year for (a) genotypic richness (*R*), (b) genotypic evenness (*D\**), (c) the distribution of clonal membership (*Pareto ß*), and (d) the multilocus estimate of linkage disequilibrium (*r̄_D_*) using Dataset Two, including all reproductive tetrasporophytes and all heterozygous thalli. Boxes represent the interquartile range, the middle lines are medians, whiskers represent the 1.5 interquartile ranges, and the black dots represent outliers. Datapoints are labeled by site number (see Table 1) and jittered along the x-axes to improve visualization.

**Figure S2.**
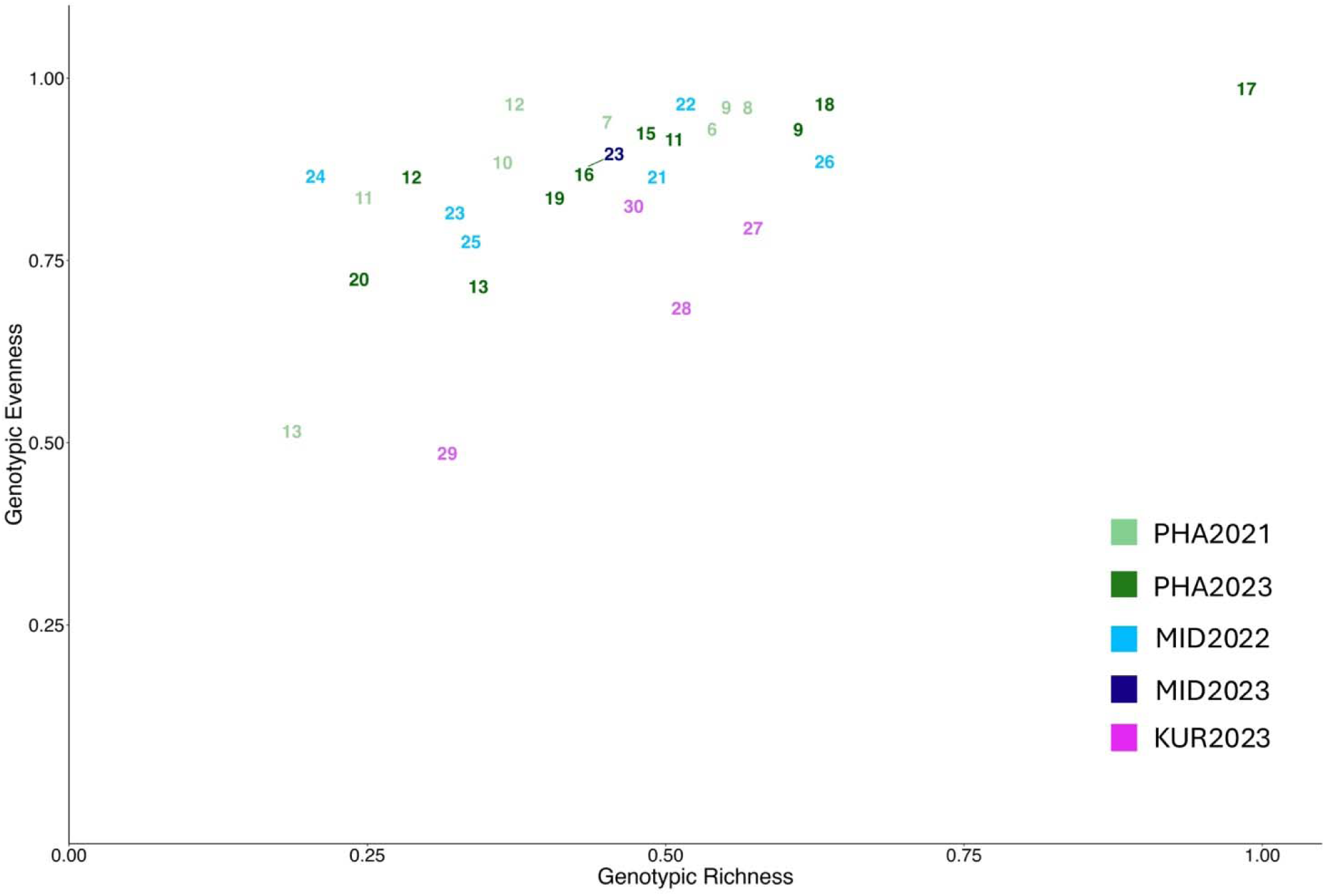
Reproductive mode variation shown as genotypic evenness (*D\**) versus genotypic richness (*R*) based on Dataset Two, including all reproductive tetrasporophytes and all heterozygous thalli. Each number refers to a site in which *Chondria tumulosa* thalli were sampled and are color coordinated by atoll and year (see Table 1). Lines are used to indicate the location of some sites on the plot to avoid overlap.

**Figure S3.**
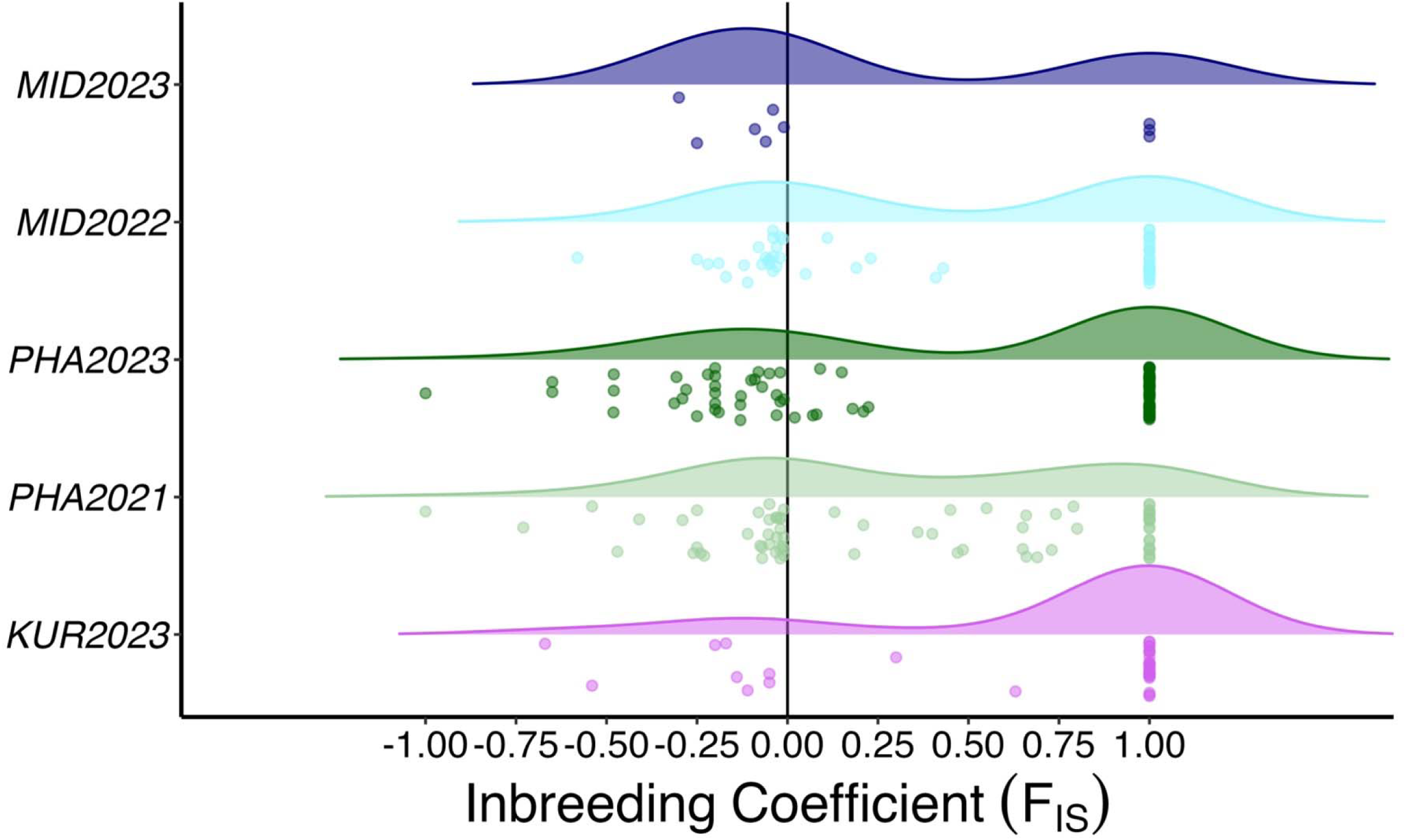
Density plot of the single locus estimates of the inbreeding coefficient (*F_IS_*) by atoll and year for sites in which we sampled *Chondria tumulosa* thalli. *F_IS_* was calculated on Dataset Two, including all reproductive tetrasporophytes and all heterozygous thalli. Number of data points representing a single locus *F_IS_* value for each site, atoll, and year are as follows: KUR2023 (36), MID2023 (9), MID2022 (54), PHA2023 (90), and PHA2021 (72).

**Figure S4.**
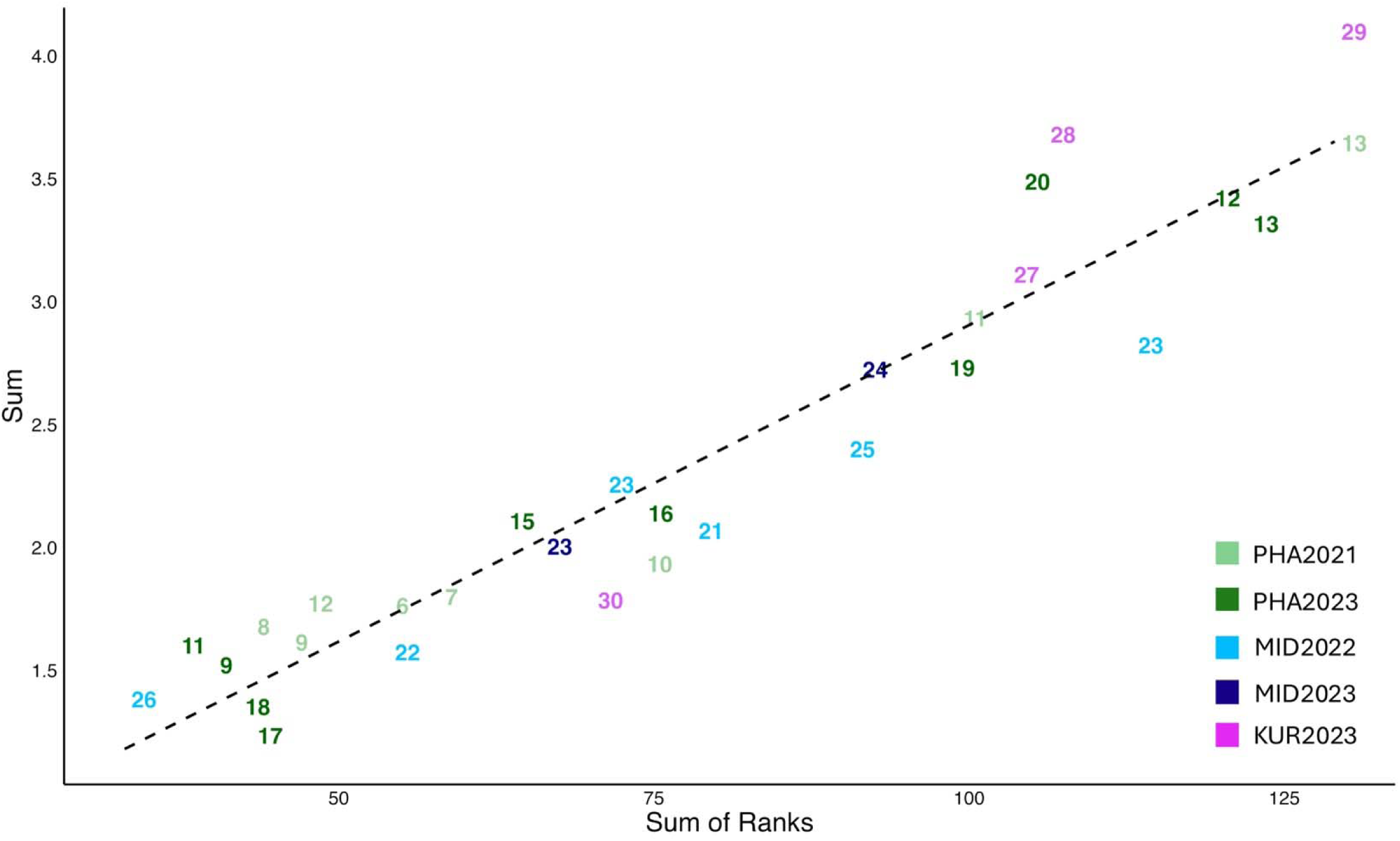
Correlation plot of the normalized and categorical ranks for all *Chondria tumulosa* sites where thalli were collected. Ranks were produced using summary statistics calculated on Dataset One including all reproductive tetrasporophytes, all heterozygous thalli, and all repeated multilocus genotypes that were identical to a reproductive tetrasporophyte (including fixed homozygous thalli). Sum of ranks is a transformed and normalized value for each site based on population genetic indices; sum is calculated by ordering each site by its population genetic index value (as described in Stoeckel et al., 2024).

**Figure S5.**
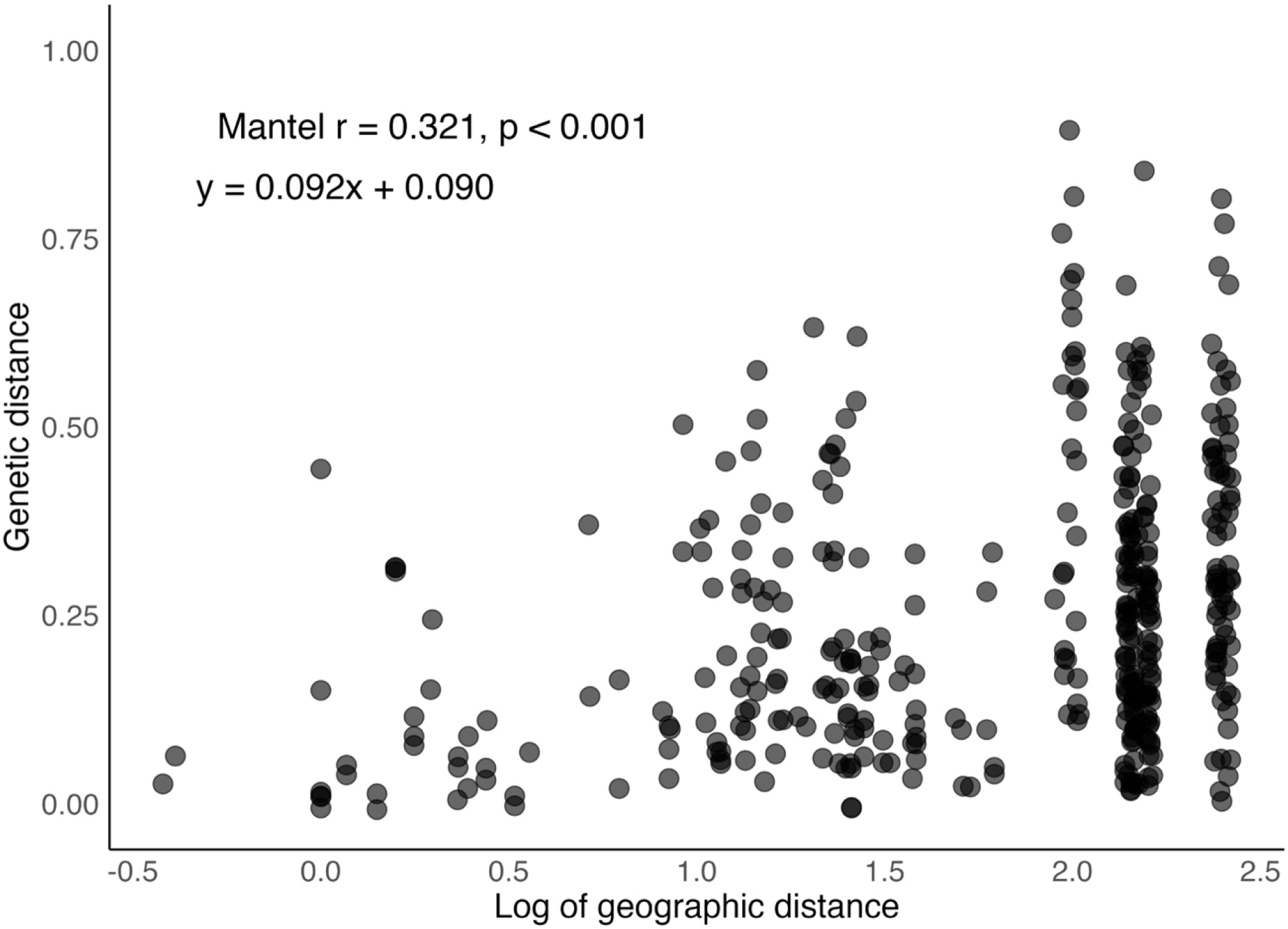
Correlation plot of indirect geographic distance and genetic differentiation for all *Chondria tumulosa* sites where thalli were collected. Genetic differentiation values were produced using Dataset One including all reproductive tetrasporophytes, all heterozygous thalli, and all repeated multilocus genotypes that were identical to a reproductive tetrasporophyte (including fixed homozygous thalli).

**Figure S6.**
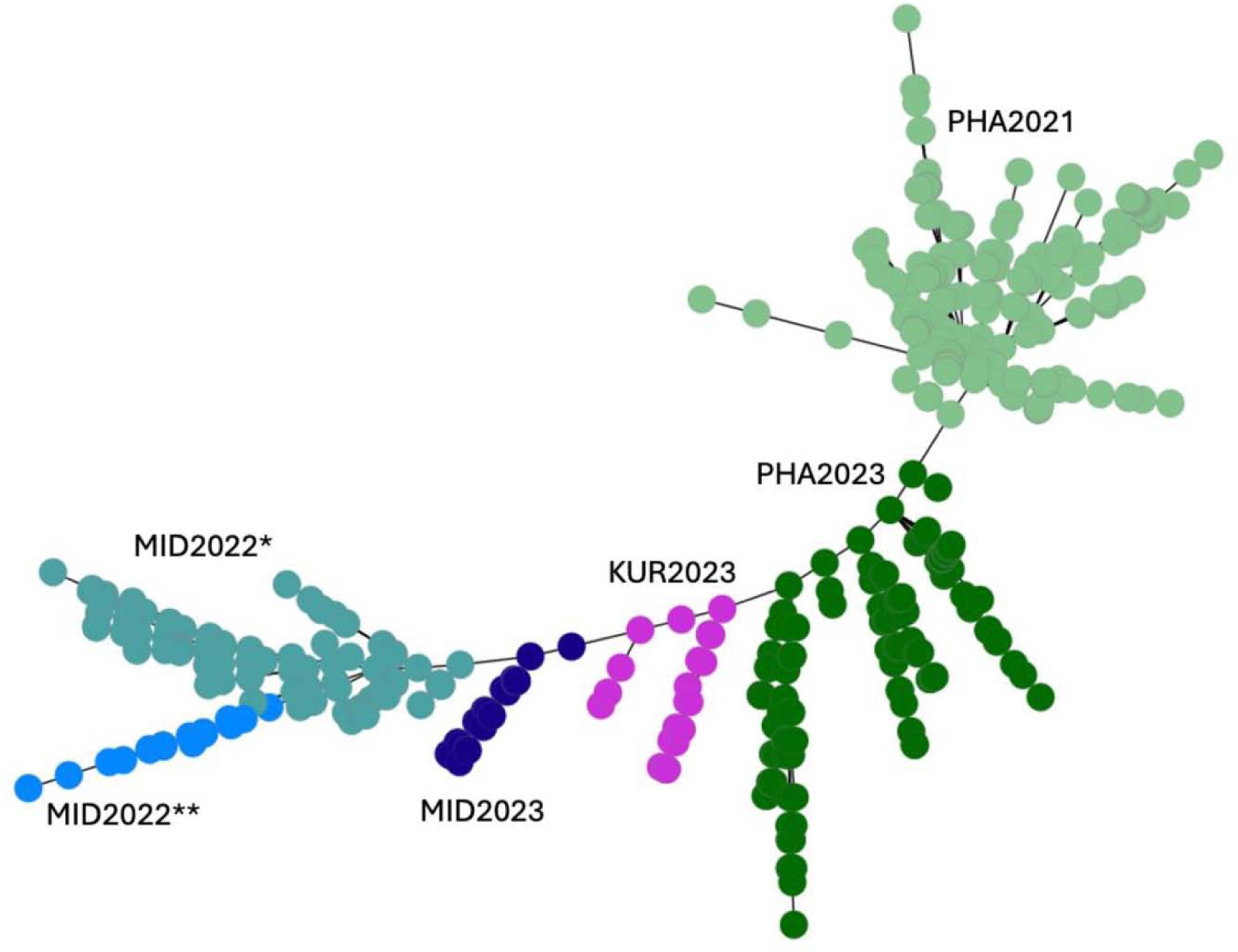
Minimum spanning tree for the sites in which we sampled *Chondria tumulosa* thalli at all locations and time points using Dataset Two, including including all reproductive tetrasporophytes and all heterozygous thalli. *Sampled in July 2022; **Sampled in October 2022.

**Figure S7.**
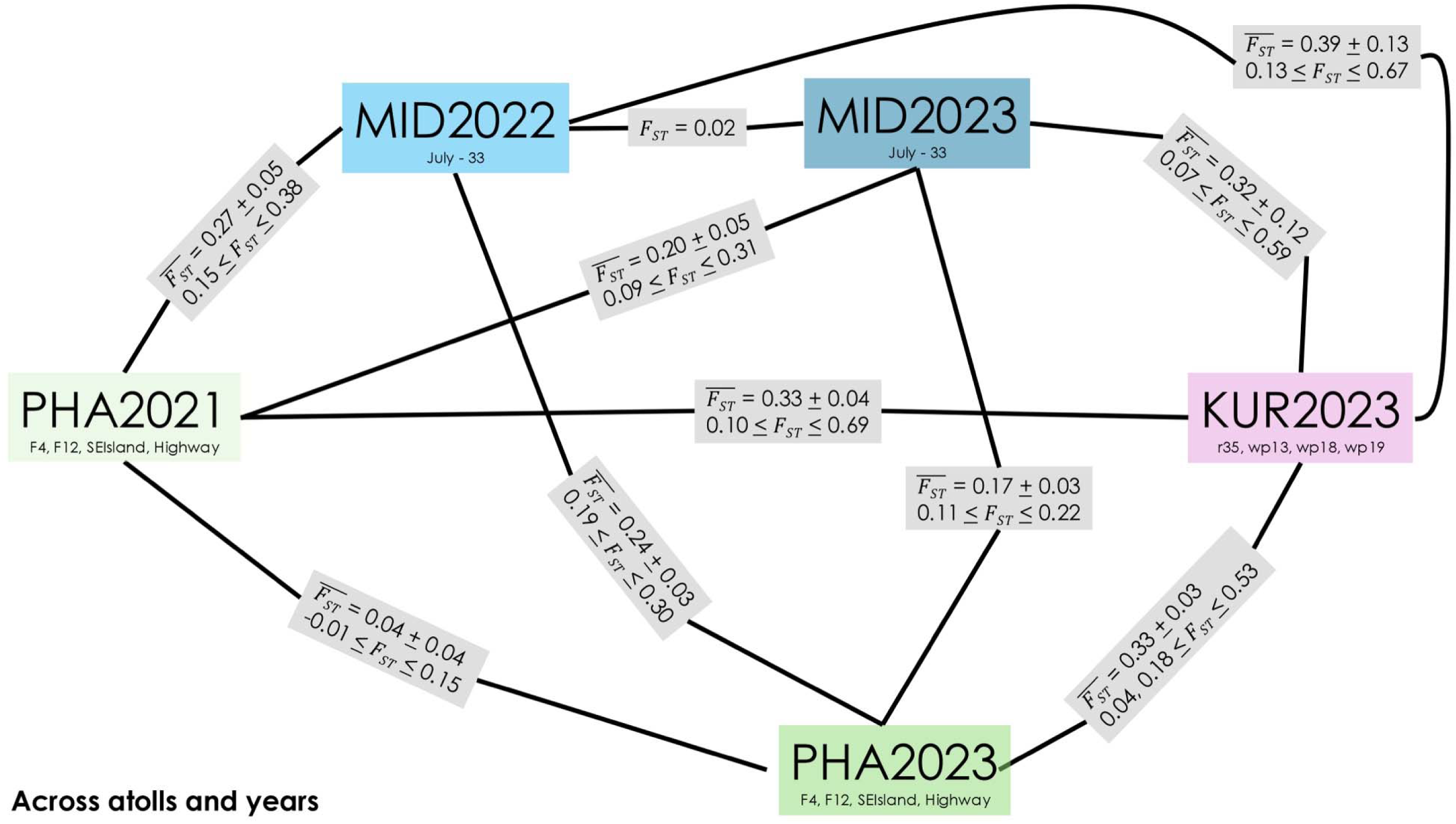
Mean *F_ST_* values from pairwise comparisons within an atoll and across atolls using only sites that *Chondria tumulosa* thalli were collected at multiple time points for Manawai (PHA_F4, PHA_F12, PHA_SEIsland, PHA_Highway) and Kuaihelani (MID_33). We used Dataset One, including all reproductive tetrasporophytes, all heterozygous thalli, and all repeated multilocus genotypes that were identical to a reproductive tetrasporophyte (including fixed homozygous thalli).

## Notes

### Competing Interest Statement

The authors have declared no competing interest.

